# Defining the structure, signals, and cellular elements of the gastric mesenchymal niche

**DOI:** 10.1101/2023.02.11.527728

**Authors:** Elisa Manieri, Guodong Tie, Davide Seruggia, Shariq Madha, Adrianna Maglieri, Kun Huang, Yuko Fujiwara, Kevin Zhang, Stuart H. Orkin, Ruiyang He, Neil McCarthy, Ramesh A. Shivdasani

## Abstract

PDGFRA-expressing mesenchyme provides a niche for intestinal stem cells. Corresponding compartments are unknown in the stomach, where corpus and antral glandular epithelia have similar niche dependencies but are structurally distinct from the intestine and from each other. Previous studies considered antrum and corpus as a whole and did not assess niche functions. Using high-resolution imaging and sequencing, we identify regional subpopulations and niche properties of purified mouse corpus and antral PDGFRA^+^ cells. PDGFRA^Hi^ sub-epithelial myofibroblasts are principal sources of BMP ligands in both gastric segments; two molecularly distinct groups distribute asymmetrically along antral glands but together fail to support epithelial organoids *in vitro*. In contrast, strategically positioned PDGFRA^Lo^ cells that express CD55 enable corpus and antral organoid growth in the absence of other cellular or soluble factors. Our study provides detailed insights into spatial, molecular, and functional organization of gastric mesenchyme and the spectrum of signaling sources for stem cell support.

## INTRODUCTION

Balanced self-renewal and differentiation of digestive epithelia relies on interactions with the underlying mesenchyme; these interactions serve as a model for adult tissue homeostasis. Stomach and intestinal epithelia share essential features, including long-lived stem cells that generate a limited number of cell types within stereotypic monoclonal crypts (intestine) or glands (stomach), but their functions and stem cell kinetics differ. *Lgr5*-expressing intestinal stem cells (ISCs) at the base of each crypt drive brisk cell turnover, whereas the gastric corpus epithelium turns over slowly, fueled by stem cells of uncertain identity and unknown molecular markers in the isthmus of each glandular unit (Figure 1A). Antral epithelium bridges the corpus and intestine, lacks villi, turns over at an intermediate rate,^1-3^ and consists mainly of foveolar and mucous cells. Antral stem cells express *Lgr5* and concentrate near the gland base (Figure 1A). Recent studies have defined anatomic and functional relations between intestinal stromal and epithelial sub-compartments.^4-8^ In contrast, gastric mesenchyme remains superficially characterized; a recent single-cell analysis combined corpus and antral cells, obscuring their individual properties.^9^ The dense stroma underlying corpus and antral epithelia contains blood vessels, smooth muscle, neuroglia, and poorly characterized “fibroblasts.” Here we report on our structural, molecular, and functional characterization of adult mouse corpus and antral mesenchyme at the level of whole tissues and single cells.

**Figure 1.**
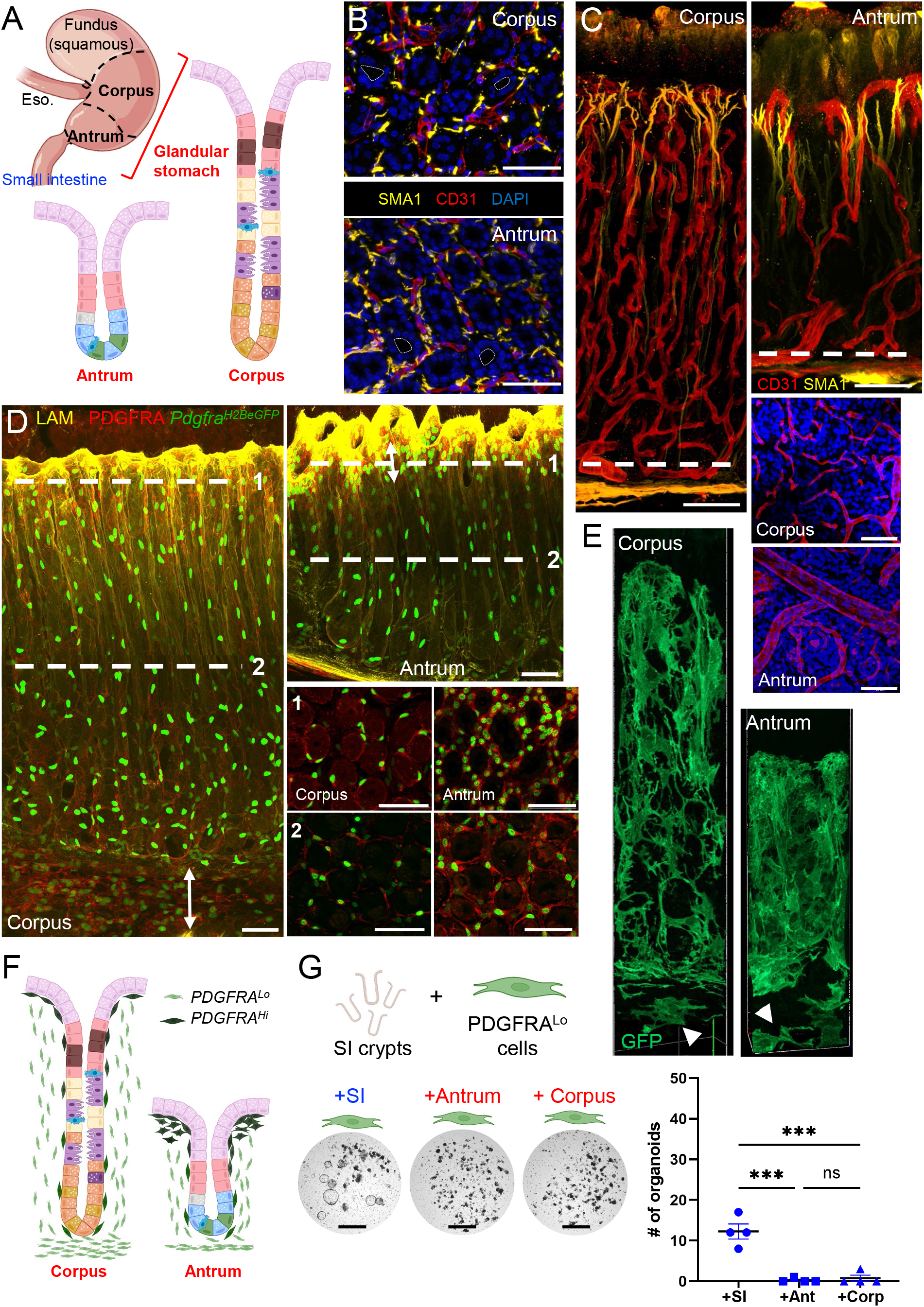
Structure and organization of gastric corpus and antral mesenchyme. **A)** Stomach and intestinal regions and different organizations of glandular stomach epithelia. **B)** Cross-sections of corpus (top) and antrum (bottom), showing smooth muscle actin (SMA1, yellow)-expressing smooth muscle fibers and CD31^+^ (red) capillaries packed tightly in the stroma between glands (nuclei stained with DAPI, blue). Dashed lines delineate gland lumen. Scale bars 50 μm. **C)** Whole-mount 3D rendering of corpus (left) and antral (right) glands, highlighting structural differences in muscle fibers (SMA1, yellow) and capillaries (CD31, red) in the two regions. Tissue cross-sections at the level marked by dashed lines are shown in the bottom right. Scale bars 50 μm. **D)** Whole-mount 3D rendering of corpus (left) and antral (top right) glands in *Pdgfra*^*H2BeGFP*^ mice. Laminin (yellow) marks the basal lamina and highlights gland pits. Tissue cross-sections at the levels marked 1 and 2 are shown in the bottom right, with PDGFRA^Hi^ cells in neon green and PDGFRA^Lo^ cells in light green. Double-headed arrow highlights PDGFRA^Lo^ layer beneath corpus gland bottoms. Immunostaining with PDGFRA antibody (red) shows the extent of PDGFRA^+^ cytoplasm enveloping each gland. Scale bars 50 μm. **E)** Single frames extracted from 3D video renderings of *Pdgfra*^*Cre(ER-T2)*^*;Rosa26*^*mT/mG*^ mouse (stained green after tamoxifen treatment) corpus (left) and antrum (right) to show the morphology of *Pdgfra*^*Hi*^ cells enveloping each gland and additional *Pdgfra*^*Lo*^ cells lying in the space beneath glands (white arrowheads). Full videos of corpus and antral gland 3D renderings are included in the Supplement, see Videos S5 and S6. **F)** Schematic representation of the distributions of PDGFRA^Hi^ (dark green) and PDGFRA^Lo^ (light green) cells in relation to corpus and antral glands. **G)** Left: Matrigel co-culture of small intestine (SI) crypts with PDGFRA^Lo^ cells isolated from SI, antral or corpus mesenchyme in the absence of recombinant factors. Unlike SI cells, stomach PDGFRA^Lo^ cells did not induce organoid formation in the absence of NOG and RSPO1. Scale bars 400 μm. Right: Quantification of organoid growth (n=4). Bars represent mean ±SEM values. Significance of differences determined by one-way ANOVA. \*\*\**p* < 0.001; ns: not significant. See also Figure S1 and Videos S1-6.

In the corpus, mucus-producing foveolar cells line a luminal “pit”; below the pit, the isthmus carries slow-cycling stem cells of uncertain identity and their transit-amplifying progeny, which produce various mature cell types (Figure 1A). The neck of each corpus gland carries mucous neck cells and acid-producing parietal cells, while pepsinogen-secreting chief cells populate the gland base. Corpus stem cell daughters migrate bidirectionally along the vertical gland axis, with foveolar precursors moving toward the pit and other precursors moving toward the gland base.^10-12^ In the antrum, proliferative epithelial cells reside in 3-4 cell tiers above the basal *Lgr5*^*+*^ stem cell zone and their progeny mainly migrate upward to replenish the physiologic attrition of terminally differentiated foveolar and mucous cells (Figure 1A).^13^ Antral stem cells require Wnt signaling to proliferate and maintain their identity,^13-15^ while bone morphogenetic protein (BMP) signaling drives epithelial differentiation^16-18^ and inhibitors of the pathway (BMPi) are implicated in gastric metaplasia and *Helicobacter pylori* infection.^19,20^ Here we define the relations between distinct digestive epithelia and their underlying stromal elements, revealing important functional differences between the small intestine (SI), gastric corpus, and gastric antrum. We identify CD55-expressing fibroblasts as the gastric equivalent of intestinal trophocytes.

## RESULTS

### Anatomic definition and signaling potential of antrum and corpus mesenchyme

Foveolar cells are present in both gastric regions; their fraction is larger in the antrum, where pits (foveolae) are wider and deeper than those in the corpus (Figure S1A). In both regions, whole-mount confocal imaging revealed blood vessels and smooth muscle fibers packed tightly in the limited space between individual glands (Figure 1B). Corpus arterioles branch along the full gland length to form a deep capillary network, while antral arterioles have a larger bore and extend capillaries mainly at gland tops, with little branching along the gland length (Figures 1C and S1B, Videos S1 and S2). Slender smooth muscle fibers extend lateral branches near the foveolar base, so the resulting fine network of lamina propria myocytes is taller and wider in the antrum (Figure 1C, Videos S3 and S4). *Pdgfra*-expressing mesenchymal cells are pivotal in intestinal epithelial homeostasis.^4^ In *Pdgfra*^*H2BeGFP*^ knock-in mice,^21,22^ we detected cells with high and low levels of nuclear GFP (Figures 1D and S1C-E). Antral PDGFRA^Hi^ cells concentrate around the pit, where they extend beyond the basement membrane into the lamina propria, while corpus PDGFRA^Hi^ cells distribute more uniformly along corpus glands (Figures S1C and S1E). PDGFRA^Lo^ cells distribute uniformly along the gland length in both segments and concentrate in layers below the gland base, at higher density in the corpus than in the antrum (Figures 1D and S1E). To assess cell morphology further, we crossed *Pdgfra*^*Cre(ER-T2)*^ knock-in mice^23^ with the *Rosa26*^*mT/mG*^ reporter strain,^24^ thus producing mice where baseline red fluorescence (TdTomato) in all cell membranes changes to green (GFP) in cells with Cre enzyme activity. Cells embedded in the basal lamina, corresponding to PDGFRA^Hi^, engulf each gland within their intercalated cytoplasmic projections and cells corresponding to PDGFRA^Lo^ are recognized beneath glands (Figure 1E, Videos S5 and S6). Thus, submucosal cell arrangements are fundamentally similar in the glandular stomach but vary in detail in the two regions, likely reflecting distinct influences on the overlying epithelia (Figure 1F).

SI crypts form organoid structures in Matrigel supplemented with EGF, Noggin (a BMPi), and RSPO1, a Wnt potentiating agent (ENR medium).^25^ As *Lgr5*^*+*^ stem cells in the antral gland base resemble those in SI crypt bottoms, the original conditions to culture antral organoids were adapted from that foundation and include an additional requirement for Wnt3a (ENRW).^13^ Selected intestinal PDGFRA^Lo^ cells, which express BMP inhibitor (BMPi) and RSPO genes, induce intestinal crypts to form spheroids *in vitro* in the absence of any supplemental factors.^4^ PDGFRA^Lo^ cells isolated by flow cytometry from each gastric segment (Figure S1F) also induced spheroids when co-cultured, without recombinant (r) factors, with glands from the same region (Figure S1G); however, PDGFRA^Lo^ cells sorted from the gastric corpus or antrum failed to support intestinal crypts *in vitro* (Figures 1G and S1G). Thus, SI and gastric stem cells have distinct cellular support requirements and PDGFRA^Lo^ cells in each mesenchyme support homologous better than heterologous epithelial stem cells.

### Regionality of gastrointestinal Pdgfra-expressing mesenchymal cells

To identify the mesenchymal basis of gastric epithelial support, we aimed first to refine factor requirements for organoid growth, deploying culture conditions that avoid carry-over of fetal bovine serum from any component. Organoids cultured in ENRW medium expressed site-specific transcripts (Figure S2A) and we examined changes in the numbers and size of antral and corpus organoids in response to modulation of factors. Corpus glands were more sensitive than antral glands to removal of any factor and, consistent with previous reports,^26^ lack of Wnt3a or RSPO1 had the largest impact on organoid formation from both sites (Figure S2B). Compared to glands cultured in ENRW, lack of rEGF or rNoggin barely compromised organoid output (Figure S2B). Cutting Wnt3a and RSPO1 supplements in half reduced corpus organoid size modestly, while antral organoids were not adversely affected; small organoids grew from both sites in the absence of rEGF (Figure 2A). Corpus organoid size was more sensitive to rNoggin depletion than antral organoids and rEGF, but not rNoggin, improved antral organoid growth independent of Wnt3a or RSPO1 (Figures 2A and S2B). These findings reinforce Wnt, BMP, and EGF signaling requirements in gastric stem cell growth, reveal specificity in each site, and imply that mesenchymal cell products may differ to match epithelial needs.

**Figure 2.**
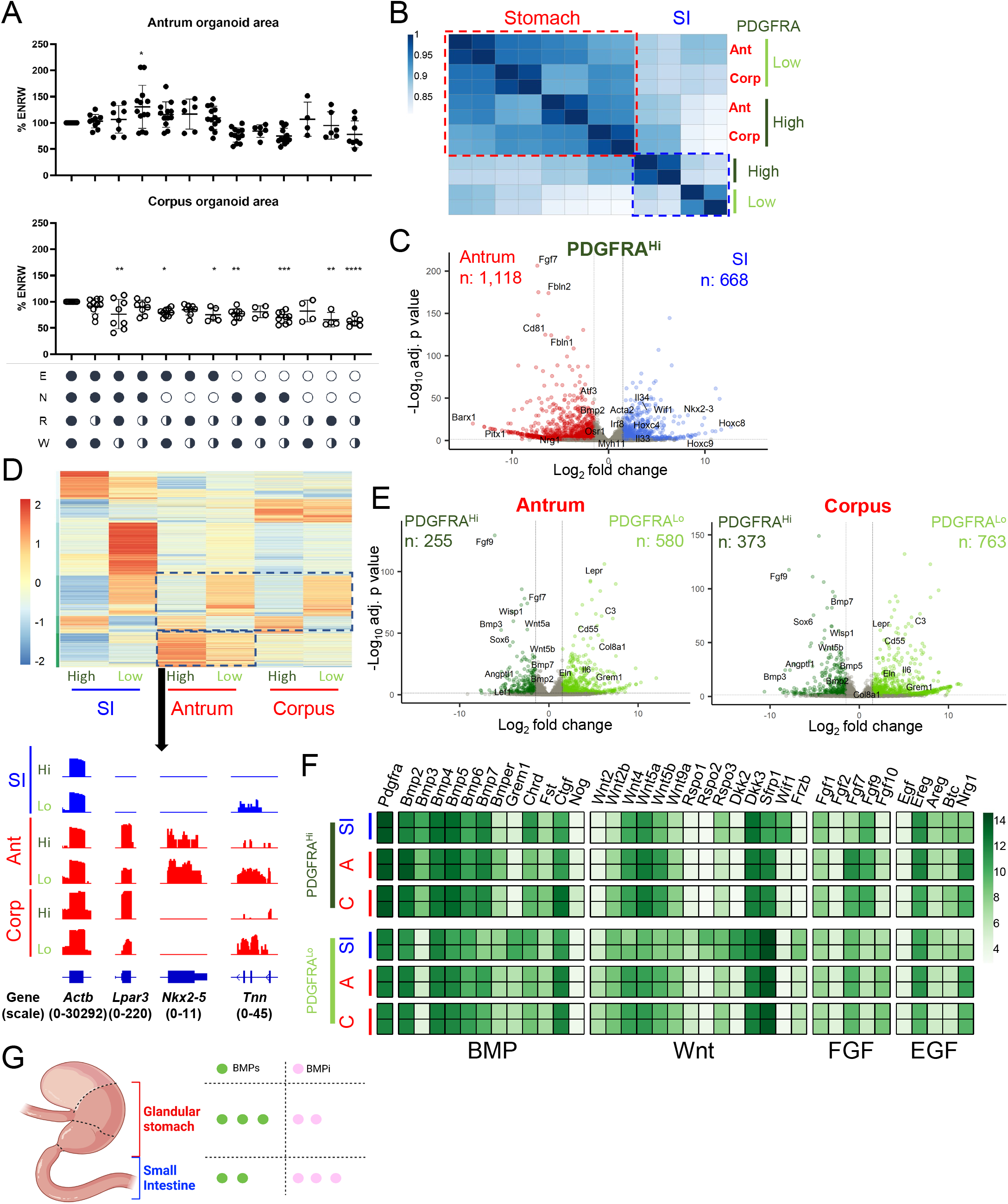
Molecular distinction of corpus and antral PDGFRA-expressing mesenchyme. **A)** Gastric organoid growth in different media, represented relative to growth in conventional ENRW medium. Circles indicate factor concentrations: full (filled circle), half, or none (empty circle). Recombinant (r) factors: E: rEGF; N: rNOG; R: medium conditioned by rRSPO1; W: medium conditioned by AFAMIN/WNT3A. Bars represent mean ±SEM values (n=4-8). Significance of differences from ENRW was determined by one-way ANOVA. \**p* <0.05; \*\**p* <0.01; \*\*\**p* < 0.001; \*\*\*\**p* <0.0001. **B)** Pearson correlations among duplicate RNA-seq libraries prepared from PDGFRA^Hi^ and PDGFRA^Lo^ cells isolated by flow cytometry after enzymatic dissociation of mesenchyme from different regions of the *Pdgfra*^*H2BeGFP*^ mouse digestive tract. SI, small intestine; Ant, antrum; Corp, corpus. Gastric PDGFRA^+^ cells are substantially different from their intestinal counterparts, as detailed in Table S1. **C)** Genes differentially expressed (*q* <0.05; log_2_ fold-difference >1.5) between antrum and small intestine (SI) PDGFRA^Hi^ cells. Selected genes are labeled. **D)** Genes enriched in each regional population (*q* <0.05; log_2_ fold-difference >1.5; *k*-means = 6) and Integrative Genome Viewer (IGV) tracks showing RNA-seq data for representative genes enriched in antral PDGFRA^+^ cells. *Actb* controls for proper normalization of read counts and the numbers in parentheses refer to the range of signal values. **E)** Genes differentially expressed (*q* <0.05; log_2_ fold-difference >1.5) between PDGFRA^Hi^ and PDGFRA^Lo^ cells in the gastric antrum (left) and corpus (right). Selected genes are labeled. **F)** Relative expression of mRNAs for known niche factors in each mesenchymal cell population, selected from the full set of BMP, Wnt, FGF, and EGF pathway agonists and antagonists. The heatmap is prepared from normalized rlog counts from DEseq2, each row represents one replicate. **G)** Graphical summary of the findings. Bulk RNA-seq data point to a higher BMP tone from *Pdgfra*^+^ cells in stomach than in the SI. See also Figure S2 and Table S1.

PDGFRA^Hi^ intestinal sub-epithelial myofibroblasts (ISEMFs) produce differentiation-inducing BMP ligands and sub-cryptal CD81^+^ PDGFRA^Lo^ “trophocytes” best oppose that signal to expand ISCs *in vitro*.^4^ Although stomach mesenchymal PDGFRA^Lo^ cells lack CD81 heterogeneity (as we show below by single-cell RNA-seq), segment-specific supportive activity of bulk PDGFRA^Lo^ cells (Figure 1G) implies that they provide certain key factors. To dissect regional mesenchymal differences, first we profiled bulk transcriptomes of PDGFRA^+^ cells purified by flow cytometry from the corpus and antrum of *Pdgfra*^*H2BeGFP*^ mice. Stripping external muscle before cell isolation excluded PDGFRA^Hi^ neurons nestled therein, so the remaining PDGFRA^Hi^ cells represent gastric SEMFs. Replicate RNA-seq libraries from each cell isolation were highly concordant (Figure S2C). In unsupervised hierarchical analysis, gastric PDGFRA^+^ cells were distinct from their previously described intestinal counterparts,^4^ followed by differences between PDGFRA^Hi^ and PDGFRA^Lo^ cells in each organ (Figure 2B). Corpus and antral PDGFRA^Hi^ cell (SEMF) transcriptomes resembled each other, compared to the substantial differences between SEMFs in the SI and those from either stomach segment (Figures 2C, S2D, and Table S1). Exclusive expression of known transcriptional regulators *Barx1* and *Pitx1* in gastric cells and of *Nkx2-3* and posterior *Hox* genes in intestinal cells support the veracity of additional findings, for example that gastric SEMFs express higher levels of *Fgf7* (a gene we examine later in this study), *Fbln1* and *Fbln2, Cd81*, and transcription factor genes *Atf3* and *Osr1*. Conversely, intestinal SEMFs express higher levels of *Il33* and *Il34*, the Wnt inhibitor *Wif1*, and transcription factor *Irf8*; differential expression of *Acta2* and *Myh11* suggests that intestinal and antral SEMFs are likely more contractile than their corpus counterparts (Figures S2E and Table S1).

Segment- and cell type-specific gene modules were evident in each population (Figure 2D), with certain elements, e.g., *Hox* genes, enriched as expected in both SEMFs and PDGFRA^Lo^ cells (Table S1). Intestinal PDGFRA^Lo^ cells have the most distinctive transcriptome of any population, with unique expression of *Eln, Col8a1*, and *Il6*, among other factors (Figure S2D). Where intestinal SEMFs and PDGFRA^Lo^ cells differ from each other at >2,100 genes, compatible with distinct and opposing functions,^4^ the corresponding populations in both stomach segments are less divergent, differing in expression of ∼800 (antrum) to ∼1,100 (corpus) genes (Figure 2E). Gastric PDGFRA^Lo^ cells from either segment are enriched for *Cd55* (a marker we investigate later in this study), *Lepr* and *Dll1* (genes that mark specific mesenchymal cells in mouse and human intestinal mesenchyme, respectively)^27,28^ while gastric SEMFs are enriched for signaling factors such as *Fgf9* and *Angptl1*, multiple BMP genes, and for transcription factor (TF) gene *Sox6* and, in the antrum, Wnt-responsive *Lef1* (Figures 2C-D, S2E, and Table S1). *Foxl1*, which marks intestinal SEMFs among other cells,^4,5,8^ is expressed in gastric SEMFs and, at a lower level, also in gastric PDGFRA^Lo^ cells. Thus, regional differences in the spatial arrangements of *Pdgfra*-expressing cells (Figure 1) extend to their gene expression profiles.

Among potential sources of the signals that support epithelial expansion *in vitro*, genes for canonical *Wnt2* and for non-canonical ligands *Wnt4, Wnt5a*, and *Wnt5b* are expressed in both PDGFRA^+^ populations in both stomach segments; however, Wnt-potentiating factors *Rspo1, Rspo2*, and *Rspo3* express at lower levels in gastric than in intestinal PDGFRA^Lo^ cells (Figure 2F). Thus, while PDGFRA^+^ cells in every segment produce Wnt ligands, only transcripts in intestinal PDGFRA^Lo^ cells display a potential for significant canonical Wnt signaling. BMP transcripts are enriched not only in intestinal, antral, and corpus PDGFRA^Hi^ cells, but *Bmp2, Bmp5*, and *Bmp7* are also prominent in gastric PDGFRA^Lo^ cells (Figure 2F). *Grem1*, a BMPi abundant in intestinal PDGFRA^Lo^ cells, and other classical BMPi are barely present in bulk populations of gastric PDGFRA^Lo^ cells, which instead express a putative and atypical BMP7 inhibitor, *Ctgf*.^29,30^ Based on mRNA levels alone, these findings suggest that overall BMP “tone” may be higher in the stomach than in the SI (Figure 2G).

Gastric organoids generally display spheroid morphology (see Figure 1G), distinct from intestinal organoids, in which budding protrusions represent crypts housing progenitor and stem cells.^25^ When culture conditions were first defined for antral organoid growth, addition of human fibroblast growth factor (FGF) 10 to the medium induced budding.^13^ Although the *in vivo* significance of this observation is unclear (glandular stomach epithelium lacks crypt structures), the asymmetry of FGF mRNA expression in PDGFRA^+^ cells is striking. Of the five transcripts expressed in any population, *Fgf9* is expressed in SEMFs and *Fgf10* in PDGFRA^Lo^ cells from all sites, while *Fgf7* is especially high in antral SEMFs. *Nrg1*, an EGF-family factor implicated in ISC regeneration after injury,^31^ is also uniquely higher in gastric (especially antral) than in intestinal SEMFs and PDGFRA^Lo^ cells.

### Transcriptional features of antral and corpus mesenchyme

Having identified substantive differences in sub-epithelial structure and signaling capacity, we examined cellular heterogeneity by mRNA profiling of whole gastric mesenchyme (manually depleted of external smooth muscle) at single-cell resolution. A total of 34,445 mesenchymal cells (20,624 from corpus and 13,821 from antrum; epithelial contamination was negligible) provided information on at least 2,000 transcripts each (see Figure S3A). Consistent with the structures and cells revealed by fluorescence microscopy, graph-based clustering identified substantial fractions of *Pecam1*^*+*^ endothelial cells, *Acta2*^*hi*^ *Myh11*^*hi*^ myocytes, *Acta2*^*lo*^ *Myh11*^*lo*^ *Rgs5*^*+*^ pericytes, and PDGFRA^*+*^ cells that lack these lineage markers (Figure 3A). Less than 10% of whole isolates were *Ptprc*^*+*^ immune cells, smaller than the 45% fraction present in whole SI mesenchyme^4^; only ∼1% of cells were glial and <1% showed signs of ongoing cell replication (Figure 3A). Despite structural differences in corpus and antral blood vessels and muscle fibers (Figure 1), transcriptomes of their constituent cells showed few regional differences. Significant cluster distinctions between corpus and antrum mapped exclusively to *Pdgfra*^+^ cells (Figures 3B and S3B), which express a broader panel of niche factors than other mesenchyme (Figure S3C). These findings justify our preceding attention on them (Figure 2) and our subsequent focus on the 5,896 antral and 6,594 corpus *Pdgfra*^+^ cells from our scRNA-seq study.

**Figure 3.**
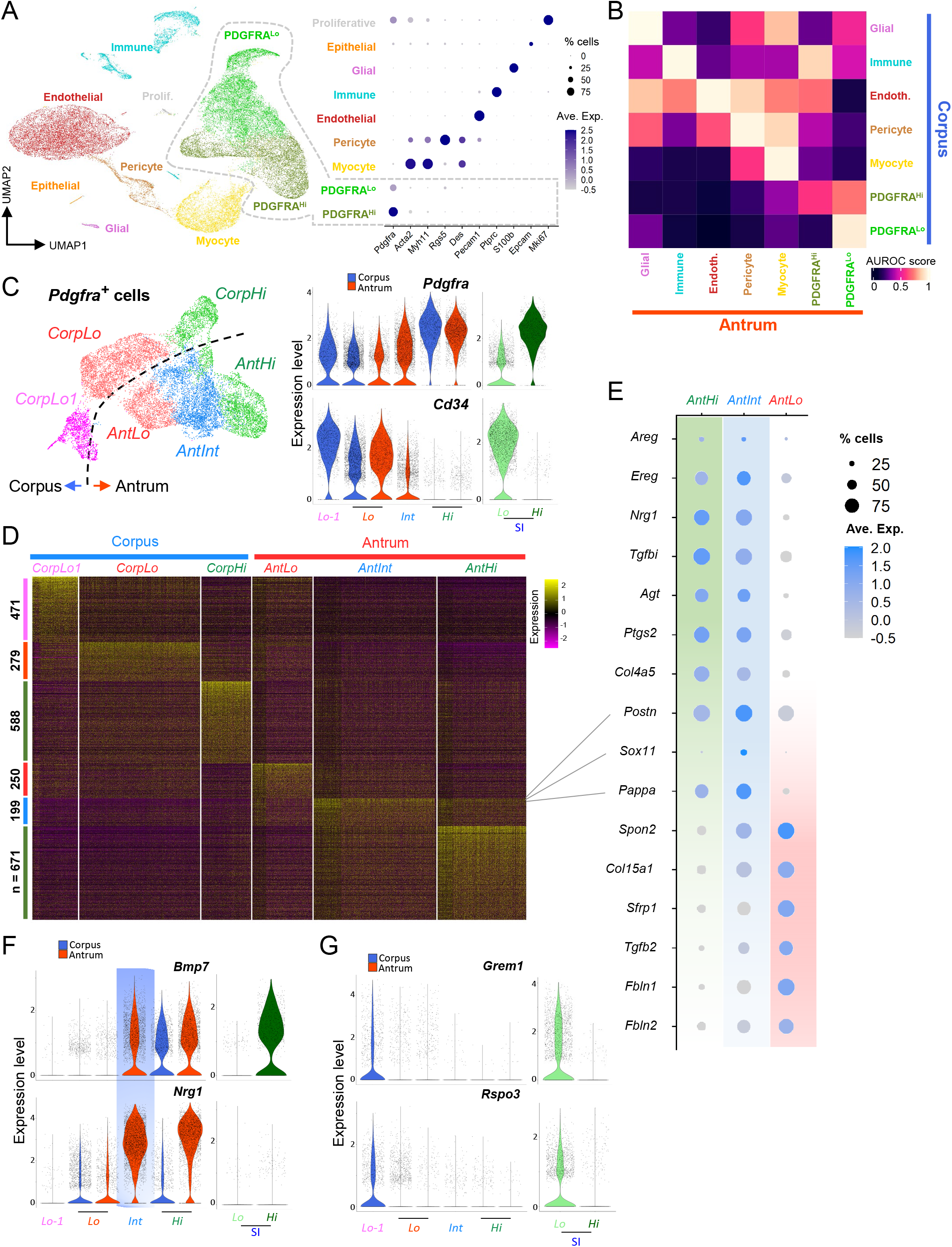
Delineation of corpus and antral PDGFRA^+^ mesenchymal cell populations. **A)** Left: Uniform manifold approximation and projection (UMAP) plot of mRNA profiles from 34,445 single cells isolated from corpus (20,624 cells) and antral (13,821 cells) mesenchyme. Right: Relative expression of known cell-restricted genes in the UMAP cell clusters. Circle diameters represent the cell fraction expressing a gene and shades of purple represent normalized average expression within the population. **B)** MetaNeighbour AUROC (area under the receiver operating characteristic) analysis shows high similarity between corresponding antral and corpus mesenchymal cells of each type other than the PDGFRA^Hi^ and PDGFRA^Lo^ cell populations. **C)** Left: UMAP identification of distinct populations of *Pdgfra*-expressing cells extracted from scRNA analysis of unfractionated mesenchyme. Right: Relative expression of *Pdgfra* and *Cd34* in the identified *Pdgfra*-expressing populations and the corresponding SI cells. Blue, corpus; red, antrum; green, SI. The 3 corpus populations are more distinct than the 3 antral populations. **D)** Differential genes expressed among the six PDGFRA^+^ mesenchymal cell types (Wilcoxon Rank Sum test, logfc.treshold = 0.25). The genes are listed in Table S2. **E)** Relative expression of marker genes from the three antral PDGFRA^+^ sub-populations identified by scRNA-seq. Circle sizes represent the percentage of cells expressing a gene, fill colors represent normalized average expression. AntInt shows distinct intermediate patterns. **F)** Expression of niche genes *Bmp7* and *Nrg1* in each population compared with *Pdgfra*-expressing SI mesenchymal cells. AntInt expression levels are comparable to those in antral *Pdgfra*^*Hi*^ cells (AntHi). *Nrg1* expression in corpus and SI is significantly lower than in antral cells. **G)** *Grem1* and *Rspo3* expression in each population compared with *Pdgfra*-expressing SI cells. Both genes are restricted to *Pdgfra*^*Lo*^ populations and expressed at lower levels in the stomach than in SI. See also Figure S3 and Table S2.

Graph-based analysis limited to these cells identified 6 clusters: *Pdgfra*^*Hi*^ and two related but distinct *Pdgfra*^*Lo*^ cell populations from each gastric segment (Figure 3C). Two *Pdgfra*^*Lo*^ cell types in the corpus, CorpLo and CorpLo1, were separate on UMAP graphs but, consistent with bulk RNA profiles (Figure 2D), antral cells separated less well. UMAP distinguished antral *Pdgfra*^*Hi*^ (AntHi) and *Pdgfra*^*Lo*^ (AntLo) cells and a substantial third pool (AntInt, 44% of *Pdgfra*^*+*^ cells) with intermediate *Pdgfra* and *Cd34* levels and a distinctive RNA profile (Figure 3C-D). That profile includes enriched expression of the transcription factor (TF) gene *Sox11*, contractile genes *Acta2* and *Myh11*, and prostaglandin synthase *Ptgs2* (Figure 3E and Table S2). Some features are similar to AntHi (e.g., EGF-family genes *Ereg, Areg and Nrg1*, the TGF-ß inhibitor *Tgfbi*, and *Col4a5*) and others to AntLo (e.g., *Tgfb2*, Wnt antagonist *Sfrp1*, and extracellular matrix factors *Spon2, Col15a1, Fbln1* and *Fbln2*). Notably, *Bmp2/5/7* and non-canonical *Wnt4* transcript levels approach those in antral and intestinal SEMFs (*Pdgfra*^*Hi*^ cells), while other signals (e.g., *Wnt5a, Fgf7*) are lower than in SEMFs (Figures 3E-F and S3D-E). Because flow cytometry separated PDGFRA^Hi^ and PDGFRA^Lo^ cells from *Pdgfra*^*H2BeGFP*^ mouse corpus better than the antrum (Figure S2A), we infer that AntInt divides between these two populations in FACS isolations. The above differences, however, mark them as distinct cells, overall resembling AntLo more than AntHi (Figure S3F), and we show below that they differ from AntLo in niche activity.

Differences between corpus and antral *Pdgfra*^*Lo*^ subpopulations also include expression of known niche factors. Low levels of intestinal niche factors *Grem1* and *Rspo3* apparent in bulk RNA profiles of stomach PDGFRA^Lo^ cells (Figure 2E) are largely limited to the distinct CorpLo1 cluster and AntLo, albeit at levels lower than intestinal *Pdgfra*^*Lo*^ cells (Figure 3G). CorpLo1 also expresses a different spectrum of extracellular matrix genes (high *Has1, Dpt* and *Col14a1* – Table S2) compared to CorpLo, suggesting that these cells may localize in different anatomic domains. In summary, we identify distinct gastric PDGFRA^+^ populations from each region and below we address their properties systematically. Of note, BMP agonist mRNA levels diverge less between *Pdgfra*^*Hi*^ and other *Pdgfra*^*+*^ cells in the stomach than in the intestine and BMPi express at lower levels (Figure S3E), confirming ostensibly higher BMP tone in the stomach.

### Molecular and spatial heterogeneity of antral SEMFs

We first considered *Pdgfra*^*Hi*^ cells, which occupy a specific anatomic compartment and have unique mRNA profiles (Figures 1-3), apart from the others. Gastric *Pdgfra*^*Hi*^ cells (SEMFs) differ from intestinal SEMFs in high *Fgf7* (antrum) and *Fgf9* (corpus) expression and in profiles of the TF genes *Foxl1* and *Gli1* (Figures S3D and S4A). Considered separately from other mesenchymal cells, they divided into one corpus and two antral subpopulations, all of which express canonical BMP ligand genes (Figures 4A and S4B). Notably, one antral cluster expresses high *Fgf7* and the EGF ligand *Nrg1*, while the other is marked by high *Bmp3* and *Ctgf* (candidate non-canonical BMPi)^32,33^ and low levels of *Nrg1*, a profile similar to corpus SEMFs (Figure 4A-B); *Fgf7* and *Bmp3* are largely SEMF-restricted, while the other genes are enriched in SEMFs but also expressed in other PDGFRA^*+*^ cells (Figures S3D-E). In situ hybridization localized antral *Fgf7*^*+*^ *Nrg1*^*+*^ SEMFs to the tops of antral glands, adjacent to foveolae, whereas *Bmp3* and *Ctgf* localize primarily in SEMFs that abut the zone of epithelial cell replication in the lower gland (Figure 4C). As antral SEMF density is higher at gland tops than at the base (Figure S1E), the proportions of SEMF subpopulations (Figure 4A) further support that molecularly distinct antral SEMF subpopulations are spatially segregated.

**Figure 4.**
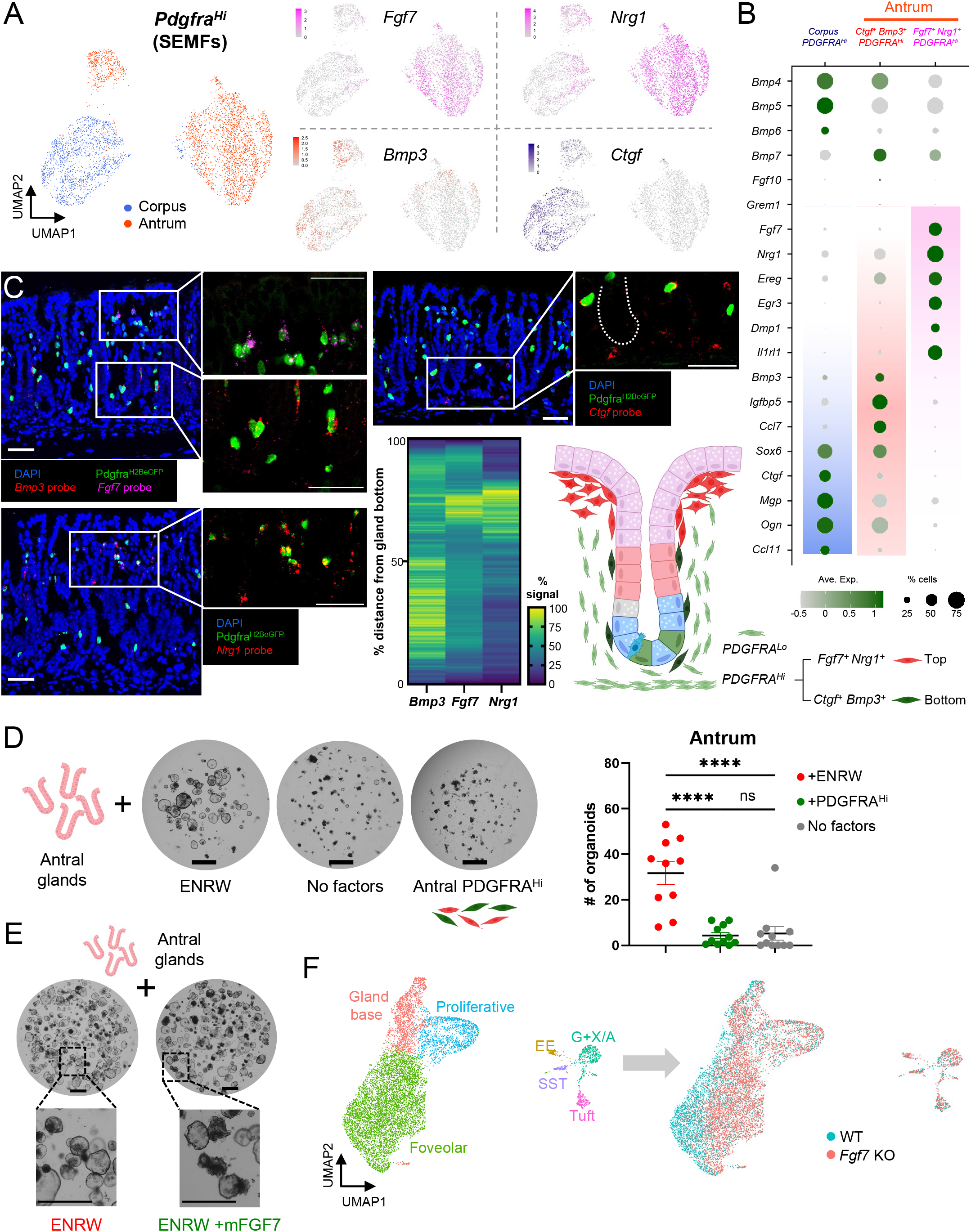
Identification of regionally distinct gastric SEMF populations and assessment of SEMF function. **A)** Left: Examined in isolation, gastric PDGFRA^Hi^ SEMFs of corpus or antral derivation are distinct and antral SEMFs divide further into two cell populations of different size. Right: Transcript density of *Fgf7, Nrg1, Bmp3*, and *Ctgf* projected onto the UMAP plot. **B)** Relative expression of marker genes from the three SEMF populations identified by scRNA-seq. Circle diameters and fill colors represent the fraction of cells expressing a gene and normalized average expression levels, respectively. The two antral SEMF populations differ in expression of *Fgf7, Nrg1, Bmp3*, and other factors; corpus SEMFs resemble the smaller antral *Bmp3*^*+*^ cluster more than the larger population of antral *Fgf7*^*+*^ *Nrg1*^*+*^ cells. **C)** Fluorescence *in situ* hybridization (RNAscope) on antral tissue sections localizes *Fgf7*- and *Nrg1*-expressing PDGFRA^Hi^ SEMFs near gland pits and *Bmp3*- and *Ctgf*-expressing cells in the lower half of glands, near epithelial stem and progenitor cells. Images represent fields examined in 3 independent experiments with each probe. Scale bars 50 μm. The heatmap displays average fluorescence signal strength quantified along 16-25 individual glands. Ctgf probe quantification is shown in Figure 5C. Right: schematic representation of the distribution of antral SEMFs. **D)** Matrigel co-culture of antral glands with unfractionated antral PDGFRA^Hi^ cells, which fail to induce organoids in the absence of rNOG and RSPO1. Glands cultured in ENRW medium serve as a control. Scale bars 400 μm. Results are quantified to the right. Bars represent mean ±SEM values from n=10 independent experiments. Significance of differences was determined by one-way ANOVA. \*\*\*\**p* <0.0001, ns: not significant. Corpus gland cultures are shown in Figure S4C. **E)** Antral glands exposed to recombinant FGF7 in addition to complete ENRW medium show an increase in budding structures, which are quantified in Figure S4D. Scale bars 400 μm. **F)** UMAP clustering of antral epithelial cells from RNA profiling of 3,885 wild-type (WT) and 5,595 mesenchymal *Fgf7*-deficient mice 90 days after *Pdgfra*^*Cre(ER-T2)*^ animals were treated with tamoxifen. Antral epithelium is essentially unchanged. Although WT and mutant foveolar cells did not overlap perfectly, differences in gene expression (listed in Table S3) were subtle. See also Figure S4 and Table S3.

Because *Bmp3* and *Ctgf* may have BMPi activity^32,33^ and SEMFs expressing those genes lie near proliferative epithelial cells, we asked if they might support antral gland growth *in vitro*; to optimize the organoid assay for BMPi activity, we reduced RSPO and Wnt3 concentrations 5-fold. In these conditions, rNOG readily supported antral organoids, but the same concentration of human rBMP3 or rat rCTGF lacked that activity (Figure S4C). No surface marker reliably separated *Bmp3*^+^ *Ctgf*^*+*^ from *Fgf7*^*+*^ *Nrg1*^*+*^ SEMFs, but unfractionated FACS-sorted antral PDGFRA^Hi^ cells (GFP^Hi^ cells from *Pdgfra*^*H2BeGFP*^ mice) did not induce organoid growth from antral glands (Figures 4D). Corpus PDGFRA^Hi^ cells similarly failed to support corpus glands *in vitro* (Figure S4D); these outcomes differ from the robust stimulation by bulk PDGFRA^Lo^ cells (Figure 1G). Together, these findings reveal spatial and molecular asymmetry of antral SEMFs without identifying supportive activity for corpus or antral stem-cell compartments, much as intestinal SEMFs lack a discernible support function in organoid assays.^4^

In antral organoids, recombinant human FGF10 induces buds, which are presumed to reflect epithelial maturation.^13^ Because *Fgf7* is abundant in peri-foveolar antral SEMFs, belongs in the same clade as *Fgf10*, and their common receptor, *Fgfr2*, is expressed in antral epithelium^34^, we considered a function for FGF7 in epithelial differentiation. Indeed, addition of murine rFGF7 to ENRW medium increased antral organoid budding substantially without affecting organoid numbers (Figures 4E and S4E). Constitutive *Fgf7* gene disruption in mice has no reported effects on gastric development or function^35^ but subtle differentiation defects could have been masked. Therefore, to assess putative gastric functions in adult mice, we used CRISPR gene editing to flank exon 2 of the *Fgf7* locus with *LoxP* sites (Figure S4F), then introduced the floxed allele onto the *Pdgfra*^*Cre(ER-T2)*^ background.^23^ *Pdgfra*^*Cre(ER-T2)*^*;Rosa26*^*LSL-tdTomato*^ mouse stomachs revealed robust Cre recombinase activity in PDGFRA^Hi^ and, at lower efficiency, in PDGFRA^Lo^ cells (Figure S4F, see also Figures 1E and Videos S5 and S6). PCR genotyping verified the expected null allele in Tomato^+^ cells isolated from antral *Pdgfra*^*Cre(ER-T2)*^*;Fgf7*^*Fl/Fl*^*;Rosa26*^*LSL-TdTomato*^ mesenchyme (Figure S4F). We detected no overt histologic defects or disturbed distribution of antral epithelial proliferative or differentiation markers (Figure S4G) and scRNA profiles of purified wild-type (WT) and *Fgf7*^*-/-*^ antral epithelium showed intact proportions and expression characteristics of proliferative and differentiated cells (Figure 4F). Cells from the gland base and proliferative compartments in WT and *Fgf7*^*-/-*^ antrum clustered together; although foveolar cell clusters were not similarly superimposed, foveolar cell transcripts in *Fgf7*^*-/-*^ mice differed only subtly from the WT (Table S3). Thus, despite the abundance and specificity of *Fgf7* expression in a spatially discrete subpopulation of antral SEMFs and its effect on organoid budding, the factor’s essential and non-redundant role in maintaining the antral epithelium is at most minor and organoid assays reveal no clear stem cell niche role for SEMFs.

### A PDGFRA^Lo^ cell population sufficient to support epithelial stem cell growth

Next we examined gastric *Pdgfra*^*+*^ cell clusters other than *Pdgfra*^*Hi*^ (Figure 3). Considered separately from SEMFs, *Pdgfra*^*Lo*^ and *Pdgfra*^*Int*^ resolved into 5 objective sub-groups within 3 clusters; the smallest and most distinct cluster corresponds to CorpLo-1 and a tiny fraction of similar antral *Pdgfra*^*Lo*^ cells, while the remaining cells cluster by corpus and antral origins (Figures 5A and S5A). Within the large corpus cluster, one sub-group of cells, CorpLo2, shares expression of a panel of niche factor genes with AntLo, while the other sub-group, CorpLo3, expresses genes (e.g., *Pappa, Sox11, Postn, Nrg1* – see Figure 3D) that mark AntInt (Figures 5A and S5B). The related populations of antral *Pdgfra*^*Int*^ and CorpLo3 selectively express BMPs, which inhibit stem cells, and non-canonical Wnts, which lack known niche function (Figures S5B-C). Conversely, factors known to constitute the ISC niche, such as *Grem1* and *Rspo3*, are selectively enriched in AntLo, CorpLo2, and the isolated corpus cluster CorpLo1, which uniquely expresses sufficient *Rspo1* to be detected by scRNA-seq (Figures S5B). Thus, scRNA analysis separates gastric *Pdgfra*^*Lo*^ cells into fractions (AntLo, CorpLo1, CorpLo2) with a potential capacity to support gastric stem cells and those that seem to lack that potential (AntInt and CorpLo3).

**Figure 5.**
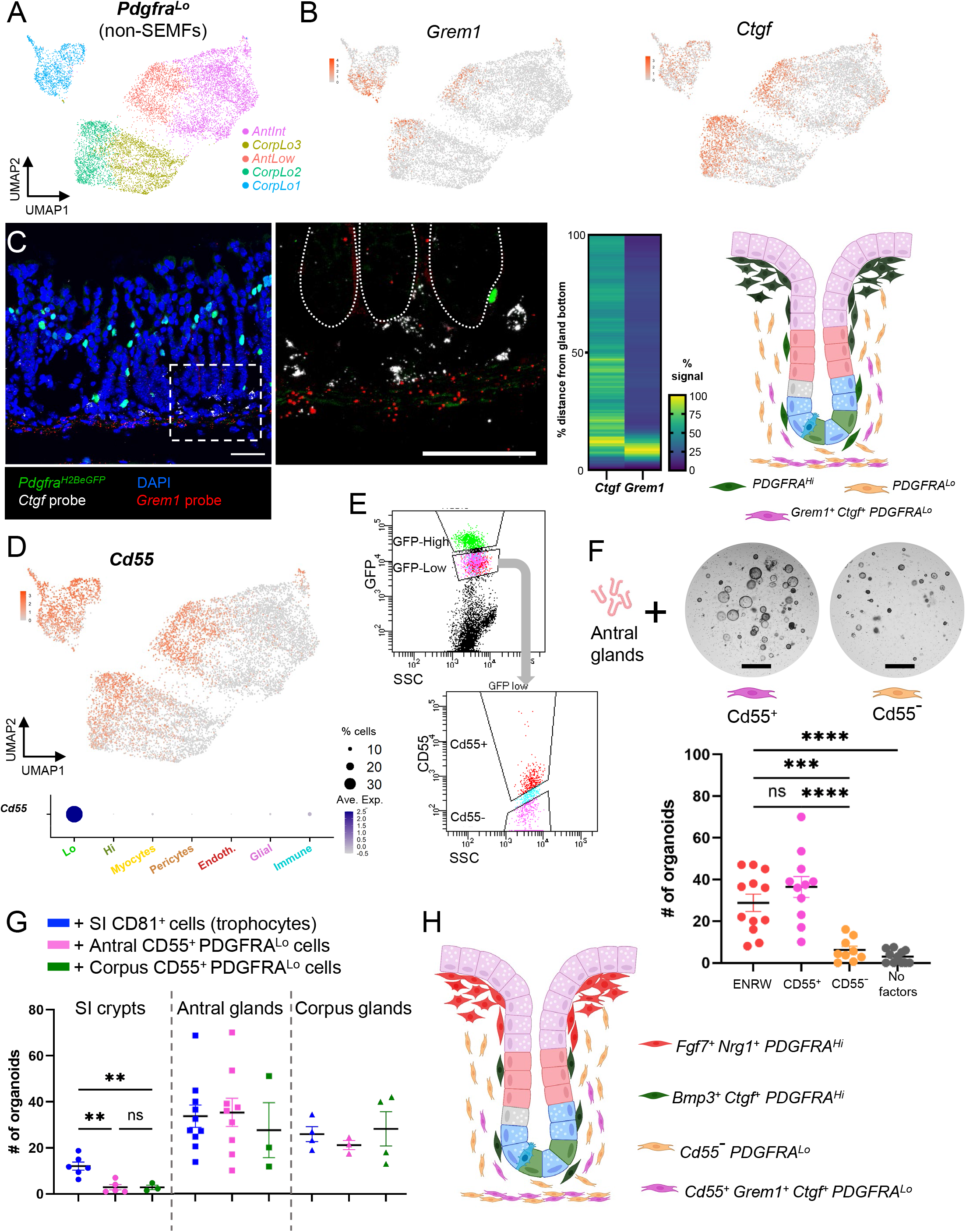
Niche activity of a CD55-expressing fraction among gastric PDGFRA^Lo^ cells. **A)** Examined in isolation, gastric PDGFRA^+^ mesenchyme other than SEMFs resolves into 5 subpopulations corresponding to the previously identified AntInt, AntLo and CorpLo1 cells (see Figure 3) and two CorpLo subpopulations: CorpLo2 and CorpLo3. **B)** Transcript density of *Grem1* and *Ctgf* projected onto the UMAP plot of these resolved cell clusters. CorpLo1, CorpLo2 and AntLo cells express the two genes almost exclusively. **C)** Left: *in situ* hybridization (RNAscope) of antral tissue sections localizes *Grem1* and *Ctgf* transcripts near the gland base, in the area corresponding to PDGFRA^Lo^ cells near the epithelial stem cell zone. Images represent fields examined in 3 independent experiments. Scale bars 50 μm. Center: Heatmap representation of fluorescence signal strength quantified along 35-39 individual glands. Right: Schematic illustration of the distribution of antral PDGFRA^Lo^ populations relative to epithelial glands. *Grem1*- and *Ctgf*-expressing cells reside selectively beneath gland bottoms. **D)** Top: Projection of *Cd55* transcript density onto the PDGFRA^Lo^ UMAP map. CorpLo1, CorpLo2 and AntLo populations show high, virtually exclusive expression. Bottom: Relative *Cd55* expression in mesenchymal and immune cells. Circle diameters and fill colors represent the fraction of cells expressing the gene and normalized average expression levels, respectively. **E)** Flow cytometry for GFP separates whole antral mesenchyme into PDGFRA^Hi^, PDGFRA^Lo^, and PDGFRA^-^ cells (top), and additional staining with CD55 antibody separates PDGFRA^Lo^ cells into CD55^+^ and CD55^-^ fractions (bottom). Isolation of the corresponding corpus population is shown in Figure S5E. **F)** Top: Matrigel co-culture of antral and corpus glands with CD55^+^ or CD55^-^ fractions of PDGFRA^Lo^ cells extracted from the mesenchyme in each gastric segment. Only the CD55^+^ fraction induces organoid formation in the absence of any recombinant factors. Scale bars 400 μm. Bottom: Organoid formation quantified from antral glands co-cultured with CD55^+^ and CD55^-^ fractions of antral PDGFRA^Lo^ cells (n= 12 independent experiments). Organoid support from PDGFRA^Lo^ CD55^+^ cells was comparable to complete ENRW medium while the CD55^-^ fraction did not induce organoid formation. Bars represent mean ±SEM values. Significance of differences was determined by one-way ANOVA. \*\*\**p* <0.001; \*\*\*\**p* <0.0001; ns, not significant. Where not indicated, differences were not significant. **G)** Quantitation of organoids formed from SI crypts and antral or corpus glands co-cultured with CD55^+^ (gastric) or CD81^+^ (SI) trophocyte fractions from each region of the digestive tract (n=3-10). Antral and corpus glands generated organoids in the presence of trophocytes from any segments, while SI crypts cultured did so only in the presence of SI trophocytes. Bars represent mean ±SEM values. Significance of differences was determined by one-way ANOVA. \*\**p* <0.01; ns: not significant. Where not indicated, differences were not significant. **H)** Schematic representation of all antral PDGFRA^+^ cell types in relation to glandular epithelium. See also Figure S5.

RNA in situ hybridization in *Pdgfra*^*H2BeGFP*^ mouse antrum localized *Grem1* and *Ctgf* in adjacent cell layers present beneath antral glands, with limited overlap between cells expressing each marker (Figure 5C). The experimental procedure largely extinguished native GFP^Lo^ cell fluorescence while preserving GFP^Hi^ signals; although the *Grem1* signal reflects expression in some *Pdgfra*^*Lo*^ cells and predominantly in the muscularis mucosae (Figure S5D, see also Figure S3C), *Ctgf* signals clearly localize to the domain of sub-glandular *Pdgfra*^*Lo*^ cells above the muscularis. In situ hybridization was unreliable in the corpus, likely owing to high mucus content. The corresponding scRNA clusters verified that *Ctgf* and *Grem1* expression domains are mutually exclusive in both regions (Figure S5E). The proximity of *Grem1*-expressing mesenchyme to stem cells at the antral gland base is reminiscent of intestinal CD81+ PDGFRA^Lo^ sub-cryptal trophocytes, which help sustain ISCs.^4^

Because unfractionated PDGFRA^Lo^ cells support antral gland growth *in vitro* in the absence of trophic factors (Figure 1G), we sought a molecular marker that might allow isolation of corpus and antral *Grem1*^*+*^ fractions for functional evaluation. *Cd55*, encoding the tetraspanin CD55, is such a gene, highly expressed in AntLo, CorpLo1, and CorpLo2, with substantially lower expression in a small fraction of immune cells (Figure 5D). To separate these PDGFRA^Lo^ mesenchymal cells away from potential immune contaminants, we performed flow cytometry with CD55 antibody on mesenchymal cells isolated from *Pdgfra*^*H2BeGFP*^ mice (Figures 5E and S5F). In co-cultures of gastric glands in basal media supplemented with CD55^+^ or Cd55^-^ fractions of PDGFRA^Lo^ cells isolated from antral or corpus mesenchyme, only CD55^+^ cells demonstrated supportive activity (Figures 5F and S5G). Neither fraction induced spheroid formation from SI crypts, but SI trophocytes induced spheroid growth from isolated corpus and antral glands (Figure 5G). Taken together, these studies reveal four distinct *Pdgfra*-expressing antral cell populations, each occupying a discrete space (Figure 5H), with niche activity selectively mapping to PDGFRA^Lo^ CD55^+^ cells that express low levels of the BMPi *Grem1* and Wnt-potentiating RSPOs; the corpus has a similar functional counterpart.

## DISCUSSION

To ensure a proper epithelial cell census, niche microenvironments around specific zones in intestinal crypts and gastric glands generate Wnt, BMP, and other signaling gradients that balance cell self-renewal against differentiation.^14,15,36-38^ Among mesenchymal cell types, PDGFRA^+^ cells as a group show the most regional diversity; in the intestine they provide canonical Wnt ligands^39^ and generate the requisite intestinal BMP gradient,^4^ but signaling sources and cellular arrangements are not known in the gastric corpus and antrum, where gland structures differ markedly.^9^ Our combination of bulk and single-cell RNA profiles, high-resolution microscopy, and organoid co-cultures identify spatially, molecularly, and functionally distinct PDGFRA^+^ cell types. Both corpus and antral mesenchyme carry three large populations, one of which expresses high *Pdgfra* and BMP and EGF ligand mRNAs, lies within the basement membrane abutting glandular epithelium, and fails to induce organoids when co-cultured with gastric glands. In each of these respects, gastric *Pdgfra*^*Hi*^ cells resemble intestinal SEMFs,^4,40-42^ which some authors call ‘telocytes’.^8,43^ A high density of *Pdgfra*^*Hi*^ cells surrounds antral gland foveolae, far from stem cells in gland bottoms, reminiscent of ISEMF aggregation at crypt and villus tops.^4,43^ Antral peri-foveolar SEMFs differ from corpus SEMFs and from deep antral SEMFs in high expression of *Fgf7, Nrg1*, and other factors that distinguish the population in nearest-neighbor analyses. We did not detect similar SEMF polarity in the gastric corpus (in part owing to technical limitations of in situ hybridization) and lone *Fgf7* gene deletion in PDGFRA^+^ cells exerted subtle consequences on pit cell gene expression; however, tight clustering of corpus SEMFs in nearest-neighbor space suggests that the population is largely uniform along the gland length. Despite the absence of a significant phenotype in our *Fgf7*-deleted mouse model, the abundance and specificity of *Fgf7* expression in peri-foveolar antral SEMFs and the budding that rFGF7 triggered in organoids suggest roles that future studies may reveal upon *Helicobacter pylori* infection or other challenges.

All other gastric PDGFRA^+^ mesenchymal cells express lower levels of *Pdgfra* and BMP and EGF ligand mRNAs, lie farther from the epithelium than SEMFs, and as a group (i.e., without further fractionation) elicit organoid formation from gastric glands in the absence of supplemental growth factors. In both gastric segments, this trophic activity maps to a distinct CD55^+^ cell fraction, which induces organoids while the complementary CD55^-^ fraction does not. In contrast to the homogeneity of corpus SEMFs, corpus *Pdgfra*^*Lo*^ cells are more heterogeneous than their antral counterparts, with the CorpLo1 pool clustering far apart from its siblings, CorpLo2 and CorpLo3 (Figure 5A). *Cd55* expression and, by extension, mesenchymal trophic activity is distributed across two corpus populations, CorpLo1 and CorpLo2, but confined to a single antral population, AntLow; the remaining antral (*Cd55*^*-*^) cells correspond to AntInt (Figure 5D). In further support of a niche function for *Cd55*^*+*^ cells, its expression is highly correlated with that of *Ctgf*, which localizes in the immediate vicinity of antral stem cells (Figure 5C – technical limitations precluded fine localization of corpus *Cd55*^*+*^/*Ctgf*^*+*^ cells by in situ hybridization). The sum of these findings leads to a simple model wherein antral *Pdgfra*^*+*^ cells other than SEMFs segregate into a niche-active CD55^+^ fraction located near epithelial stem cells and inactive AntInt, which lack CD55. Corpus CD55^+^ cells also display niche activity *in vitro*, but the precise locations and distinctions among corpus *Pdgfra*^*+*^ cells *in vivo* are still unclear.

Although AntLo express less *Grem1* and other trophic mRNAs than intestinal trophocytes, the anatomic and functional parallels with the SI stem-cell niche are striking. Trophocytes were originally defined as CD81^+^ PDGFRA^Lo^ intestinal cells that lie beneath the muscularis mucosae and robustly support organoid formation from isolated intestinal crypts.^4,44^ Subsequently we found that supportive niche activity extends above the muscularis mucosae, into peri-cryptal CD81^-^ CD55^+^ PDGFRA^Lo^ cells.^39^ Thus, the common features of antral and intestinal cells with demonstrable niche activity are high CD55 and low PDGFRA expression, absent to low levels of BMP ligand genes *Bmp2/5/7*, and localization near epithelial stem cells. Although CD55 function in this context is unknown, its utility as a surface marker of a uniquely functional cell population is robust, supplanting the value of CD81, which is useful only in conjunction with PDGFRA. In the interest of uniformity and simplicity, we propose to redefine ‘trophocytes’ as the niche-active CD55^+^ mesenchymal cell fraction in both intestine and antrum; pending additional information on corpus CD55^+^ cell localization, possibly near the gland isthmus, the same definition may also apply in that segment. Notably, gastric CD55^+^ cells from either segment supported organoid formation from homologous or heterologous glands but not from SI crypts; this may be because their levels of *Rspo3* and *Grem1*, essential trophic factors for ISCs, are lower than those in intestinal trophocytes. The functions of CD55^-^ PDGFRA^Lo^ cell fractions, including AntInt, remain unknown in any region of the digestive tract.

In summary, we report spatial, molecular, and functional features of gastric PDGFRA^+^ mesenchyme at high resolution, noting salient differences between gastrointestinal regions. Our findings identify CD55 as a robust surface marker of PDGFRA^Lo^ cells with demonstrable niche activity, distinct from spatially segregated and functionally inert CD55^-^ cell, and they establish a foundation for further investigation of gastric mesenchymal niches in homeostasis and disease.

## Author Contributions

E.M., N.M. and R.A.S. conceived and designed the study; E.M., G.T., N.M. and A.M. performed experiments; E.M., S.M. and R.H. performed computational analyses; D.S., K.Z., Y.F. and S.H.O. generated the *Fgf7*^*fl/fl*^ mouse model; K.H. assisted with image signal quantitation; E.M. and R.A.S. analyzed and interpreted data and wrote the manuscript, with input and approval from all authors.

## Acknowledgments

Supported by NIH grant R01DK121540, the Intestinal Stem Cell Consortium of NIDDK and NIAID (U01DK103152), NIH fellowship K01DK125639, and a gift from the Sarah Rhodes Fund. We gratefully acknowledge the contributions of Molecular Imaging Core (MIC), flow cytometry, and organoid core facilities at the Harvard Digestive Diseases Center (P30DK034854) and the Dana-Farber Cancer Institute.

## STAR METHODS

### RESOURCE AVAILABILITY

#### Lead contact

Further information and requests for resources and reagents should be directed to and will be fulfilled by the lead contact.

#### Materials availability

Materials used in this study are listed in the Key Resources Table.

#### Data and code availability

Sequencing data have been deposited in the Gene Expression Omnibus (GEO) repository under the accession numbers listed in the Key Resources Table. Data will be publicly available as of the date of publication.

This study includes analysis of previously published data, referenced with the accession numbers in the Key Resources Table.

Any additional information required to reanalyze data reported in this paper is available from the lead contact upon request.

## EXPERIMENTAL MODEL DETAILS

### Mouse models and treatment

All animal procedures were approved by Animal Care and Use Committees at the Dana-Farber Cancer Institute. Mice were housed in a pathogen-free animal facility and maintained on a 12-h light/dark cycle at constant temperature and humidity, with *ad libitum* access to food and water. Strains were maintained on a mixed *C57BL/6* background. Male and females aged 8-16 weeks were used. Genotypes were identified by PCR analysis of genomic DNA isolated. *Pdgfra*^*H2BeGFP*^ (JAX strain 007669), *Pdgfra*^*Cre(ER-T2)*^ (JAX strain 032770), *Rosa26*^*Lsl-TdTomato*^ (JAX strain 007908), and *Rosa26*^*mT/mG*^ (JAX strain 07676) mouse strains were purchased from Jackson Laboratories. *Rosa26*^*mT/mG*^ mice were crossed with *Pdgfra*^*Cre(ER-T2)*^ to generate compound heterozygotes. Cre recombinase activity was induced by intra-peritoneal (i.p.) injection of 2 mg tamoxifen (Sigma Aldrich T5648) on 4 consecutive days, followed by visualization of *Pdgfra*-expressing cells. *Fgf7*^*fl/fl*^ mice, generated as described below, were crossed with *Pdgfra*^*Cre(ER-T2)*^ and *Rosa26*^*Lsl-TdTomato*^ to obtain *Pdgfra*^*Cre(ER-T2)*^*;Fgf7*^*fl/fl*^;*Rosa26*^*LSL-TdTomato*^ mice, and tamoxifen 2 mg/day was injected i.p. to induce cell-specific Cre activation and genetic recombination. *Cre*-negative littermates were used as controls.

### Fgf7 conditional mouse generation

A floxed conditional *Fgf7* allele was generated by genome editing in 1-cell embryos. The targeting vector consisted of the *Fgf7* exon 2 sequence flanked by LoxP sites and surrounded by short homology arms. Adeno-associated virus (AAVs) and electroporation were used to deliver this vector along with Cas9 protein and two sgRNA, as previously described.^45^ To prevent re-cutting after homologous recombination, single-nucleotide substitutions at the PAM motifs were included in the targeting vector. Oligonucleotides used for *in vitro* transcription of sgRNAs and for genotyping are listed in Table S4.

## METHOD DETAILS

### Gastrointestinal organoid culture

Organoids were generated from wild-type mouse SI crypts or gastric glands. Crypts were isolated from the first half of the SI by incubating the tissue at 4°C in cold 2 mM EDTA-PBS, as previously described.^25^ Crypts were plated in 5 _μ_L Matrigel (Corning 356234) droplets and cultured in DMEM/F12 medium supplemented with 1X Glutamax, 10 mM HEPES, 1X penicillin/streptomycin, 1X normocin (Invivogen, ANT-NR-2), 1X primocin (Invivogen, ANT-PM-2), 1X N2 and B27 supplements (Life Technologies, 17502001 and 17504001), 1 mM N-acetylcysteine, 50 ng/mL murine rEGF (Peprotech, 315-09), 100 ng/mL murine rNOGGIN (Peprotech, 120-10C), 10% Rspo1 conditioned medium (from 293T cells expressing mouse RSPO1). Gastric glands were isolated from the gastric antrum or corpus by incubating the tissue in 10 mM EDTA-PBS at room temperature, as described previously.^46,47^ Glands were separated from the stroma and muscularis by gentle scraping, washed in PBS, and cultured in the same medium as SI crypts with addition of 10% Afamin/Wnt3a conditioned medium (MBL International, J2-001) or as specified. Recombinant factors (murine FGF7, Peprotech, 450-60; human BMP3, Peprotech, 120-24B; rat CTGF, R&D Systems, 92-37C-T050) were resuspended in 0.1% BSA/PBS and used at final concentration 100 ng/mL.

### Quantitation of mRNA expression in organoids

After 4 days of organoid culture, Matrigel droplets were washed in cold PBS, detached gently from the plate, and incubated 20 min in cell recovery solution (Corning, 354270) to depolymerize the matrix. Organoids were collected by centrifugation, resuspended in Trizol (Thermo Fisher Scientific), and total RNA was isolated with RNeasy Mini Kit (QIAGEN). 1 _μ_g RNA was reverse transcribed using SuperScript III First-Strand Synthesis System (Invitrogen, 18080-051). Transcript levels were measured using Power SYBR Green (Life Technologies, 4367659) on a LightCycler 480 instrument (Roche) and normalized to *Gapdh* levels (2^-ΔCt^). Primer sequences are listed in Table S4.

### Mesenchymal cell isolation

SI, antral, and corpus mesenchyme was isolated as in previous reports.^4,21,48^ Tissue was harvested after perfusing adult *Pdgfra*^*H2BeGFP*^ mice with cold PBS. After manual removal of external muscles, the epithelium was removed by rotating the tissue at 250 rpm in pre-warmed chelation buffer (10 mM EDTA, 5% FBS, 1 mM Dithiothreitol-DTT, 10 mM HEPES in Hank’s Balanced Salt Solution, HBSS) for 20 min at 37°C. The tissue was then washed with PBS and digested for 10 min at 37°C in 3 mg/ml collagenase II (Worthington, LS004176) in HBSS supplemented with 5% FBS and 10 mM HEPES. Cells were harvested by centrifugation at 300*g* for 5 min. For bulk RNA sequencing, cells were sorted on a BD FACSAria II cell sorter, gating on DAPI negative cells to identify live cells, and GFP to sort *Pdgfra*-high or -low expressing cells. All DAPI-negative viable cells were collected for single cell-RNA sequencing.

### Organoid and mesenchymal cell co-cultures

Whole mesenchyme was isolated and plated on tissue culture plates (SI) or fibronectin-coated tissue culture plates (gastric mesenchyme, Corning, 354451) in DMEM/F12 medium supplemented with 10% FBS, 1X penicillin/streptomycin, 1X normocin, 1X primocin, 1X Glutamax, and 10 mM HEPES. Three days later, cells were washed in PBS, removed from the plates with 0.25% Trypsin (5 min at 37°C), washed, and stained with antibodies. SI mesenchyme was stained with biotin-CD81 antibody (eBioscience, 13081181, 1:100) followed by incubation with streptavidin-APC conjugated secondary antibody (eBioscience, 17431782, 1:100). Gastric cells were incubated with PE-conjugated CD55 antibody (BioLegend, 131803, 1:75). 5,000 viable FACS-sorted cells were co-cultured with ∼20 glands or crypts in 5 _μ_l Matrigel droplets in basal media (DMEM/F12 supplemented with 1X Glutamax, 10 mM HEPES, 1X penicillin/streptomycin, 1X normocin, 1X primocin, 1X N2 and B27 supplements, and 1 mM N-acetylcysteine with no recombinant factors).

### Immunohistochemistry and imaging

Immunohistochemistry was performed on 10 _μ_m fixed tissue sections prepared using a Leica cryostat on tissues frozen in OCT compound (Tissue-Tek, VWR Scientific, 4583). Whole-mount immunohistochemistry was performed as described.^49^ Mice were perfused with ice cold PBS, followed by 4% paraformaldehyde (PFA). Antrum or corpus was pinned on agarose plates, treated with mucolytic solution (20 mM N-acetyl cysteine and 20 mM DTT in PBS) for 15 min at room temperature, and fixed for 15 min in 4% PFA. External muscle was removed manually before overnight fixation with 15% picric acid, 0.5% PFA in PBS. On day 2, tissues were washed in PBS and placed in 10% sucrose for 4 h, then 20% sucrose overnight at 4°C. On day 3, tissues were placed in blocking buffer (0.5% bovine serum albumin, 0.6% Triton X-100, 0.05% goat serum, 0.0005% sodium azide) for 6 h, followed by overnight incubation with primary antibodies at 4°C. On day 4, after 5 hourly washes in wash buffer (0.3% Triton X-100 in PBS), tissues were incubated at 4°C with Alexa Fluor-conjugated secondary antibodies (Invitrogen) and DAPI (1:1000 dilution). The next day tissues were washed for 5 h with buffer changes every 30 min and fixed in 4% PFA at 4°C for 1-2 days. Hand-cut 1 mm fragments were positioned on glass slides with imaging spacers (Grace Bio-Labs SecureSeal, Sigma Aldrich), cleared for 30 min with FocusClear (CelExplorer, FC-101) at room temperature, and mounted with VectaShield (Thermo Fisher Scientific) medium. Tissues from at least 3 independent mice were imaged using an SP5X laser scanning confocal microscope (Leica Microsystems CMS GmbH) or an LSM980 microscope and ZEN Desk 3.4 software (Carl Zeiss GmbH). Images were analyzed using Fiji^50^ and videos were generated with ZEN Desk 3.4 software. The following primary antibodies were used at 1:500 dilution unless otherwise specified: CD31 (BD Cell Analysis, 557355); Laminin (Sigma Aldrich, L9393); alpha-smooth muscle actin1 (SMA1, Abcam, ab5694); PDGFRA (R&D Systems, AF1062, 1:100); MUC5AC (Cell Signaling Technology, 6119S); Gastrin (Abcam, ab232775, 1:100); and Somatostatin (Santa Cruz Biotechnology, sc-55565, 1:100).

### Bulk RNA sequencing

*Pdgfra*-expressing cells were isolated and FACS-sorted from antrum and corpus as described above and placed in Trizol Reagent (Thermo Fisher Scientific). Total RNA was isolated with RNeasy Micro Kit (QIAGEN) and used for library preparation (5-10 ng) using SMART-Seq v4 Ultra Low Input RNA Kit (Clontech). Libraries were sequenced on Novaseq platform (Illumina).

### Single-cell RNA sequencing

Whole mesenchyme was isolated from antrum and corpus as described above, with elimination of dead cells by flow cytometry with DAPI stain. About 10,000 cells were loaded onto a 10X Genomics Chip G. Libraries were prepared according to the manufacturer’s protocol using 10X Single-Cell 3’ v3.1 chemistry (10X Genomics, PN-1000128) and sequenced on the NovaSeq platform (Illumina).

### RNAscope in situ hybridization

mRNA detection for spatial localization of cell populations was carried out using the RNAscope Multiplex Fluorescent Reagent Kit v2 (Advanced Cell Diagnostics).^4,51^ Probes designed by Advanced Cell Diagnostic were used to detect *Grem1* (314741), *Ctgf* (314541), *Fgf7* (443521), *Bmp3* (428461), and *Nrg1* (468841) transcripts following the manufacturer’s recommended procedure after 5 min of target retrieval. GFP signal was revealed by immunohistochemistry using GFP antibody (Abcam, ab6556, 1:100) as described previously.^4^ Tissues were imaged using an SP5X laser scanning confocal microscope (Leica Microsystems CMS GmbH). Fluorescent signal distribution along gastric glands was quantified with Fiji software.

### Data processing and analysis

#### Image segmentation and quantitation

Organoid images were segmented and GFP^+^ cells were quantified using Cellpose plugin in CellProfiler.^52,53^ Four to 12 organoid images were taken into consideration per condition. Segmentation masks were generated using Cellpose pre-trained model “cyto” and a cell diameter of 150 pixels (minimum size 15 pixels, flow threshold 0.7). For GFP^+^ cells, 6 images were quantified per gastric region and representative examples are reported. Masks were generated using Cellpose pre-trained model “cyto2” and a cell diameter of 43 pixels. Each image contained about 300 segmented GFP^+^ cells and >99% of cells were included in the segmentation. Segmented masks and original images were then fed into CellProfiler to quantify organoid numbers, area occupied per organoid, and the distributions of GFP^+^ cells.

#### Bulk RNA-sequencing

Data were aligned to mouse reference genome mm10 using the Viper pipeline with default settings.^54^ Data quality was verified using RSeQC^55^ and R platform 4.2.1 was used for further analyses. Reads from genes known to express in gastric epithelium and genes encoded on sex chromosomes were excluded. Data were normalized and differential gene expression (*p*_*adj*_ <0.05; |log_2_ fold-change| >1.5) was analyzed with the DESeq2 package.^56^ Reads per kb per million tags (RPKM) values were generated from normalized counts. Pearson correlation coefficients were calculated from DESeq2 normalized counts and plotted using Corrplot package.^57^ Principal component analysis (PCA) was done and heatmaps of selected genes were generated using Rlog-transformed counts. Integrative Genome Viewer (IGV) data tracks were generated from RPKM-normalized bigwigs loaded into IGV 2.15.4 (Broad Institute). Data from SI mesenchyme (GSE130681) were reported previously.^4^

#### Single cell RNA sequencing

Samples were aligned to the mm10 genome using 10x Genomics Cell Ranger 3.0.2 with default parameters.^58^ Bioinformatic analyses were conducted using R version 4.2.1. Single-cell analyses used Seurat 4.1.0.^59^ Cells were filtered for <12% mitochondrial and >1,000 unique reads. Data were merged and normalized using sctransform.^60^ Cell features were detected with the FindVariableFeatures function (2,000 features, selection method “vst*”*). The first 15 principal components were taken to run uniform manifold approximation and projection (UMAP) algorithms.^61^ Cell types were identified using established marker genes detected by the “FindAllMarkers” function in Seurat2 (min.pct=0.25, test “roc”).

MetaNeighbor-AUROC analysis, a robust tool to assess similarities between cell types, was used to compare gastric mesenchymal populations. In brief, highly variable genes identified in the datasets were used to train an algorithm to predict the type of a given cell by the expression of its nearest neighbors. To calculate ‘similarity’ between cell types A and B across two samples, the cell type annotation from one group was hidden and a score was used to represents how often a cell of type A is predicted to overlap with cells of type B versus all other cell types (area under the receiver operating characteristic, AUROC score). Higher scores indicate greater similarity between two cell types.

## QUANTIFICATION AND STATISTICAL ANALYSIS

Prism9 (GraphPad Inc) was used for statistical analysis of all data. Specific tests and significance levels are indicated in figure legends. Graphical drawings were created at BioRender.com

