## Supplemental Figures for "Defining the structure, signals, and cellular elements of the gastric mesenchymal niche"

Manieri et al, Figure S1

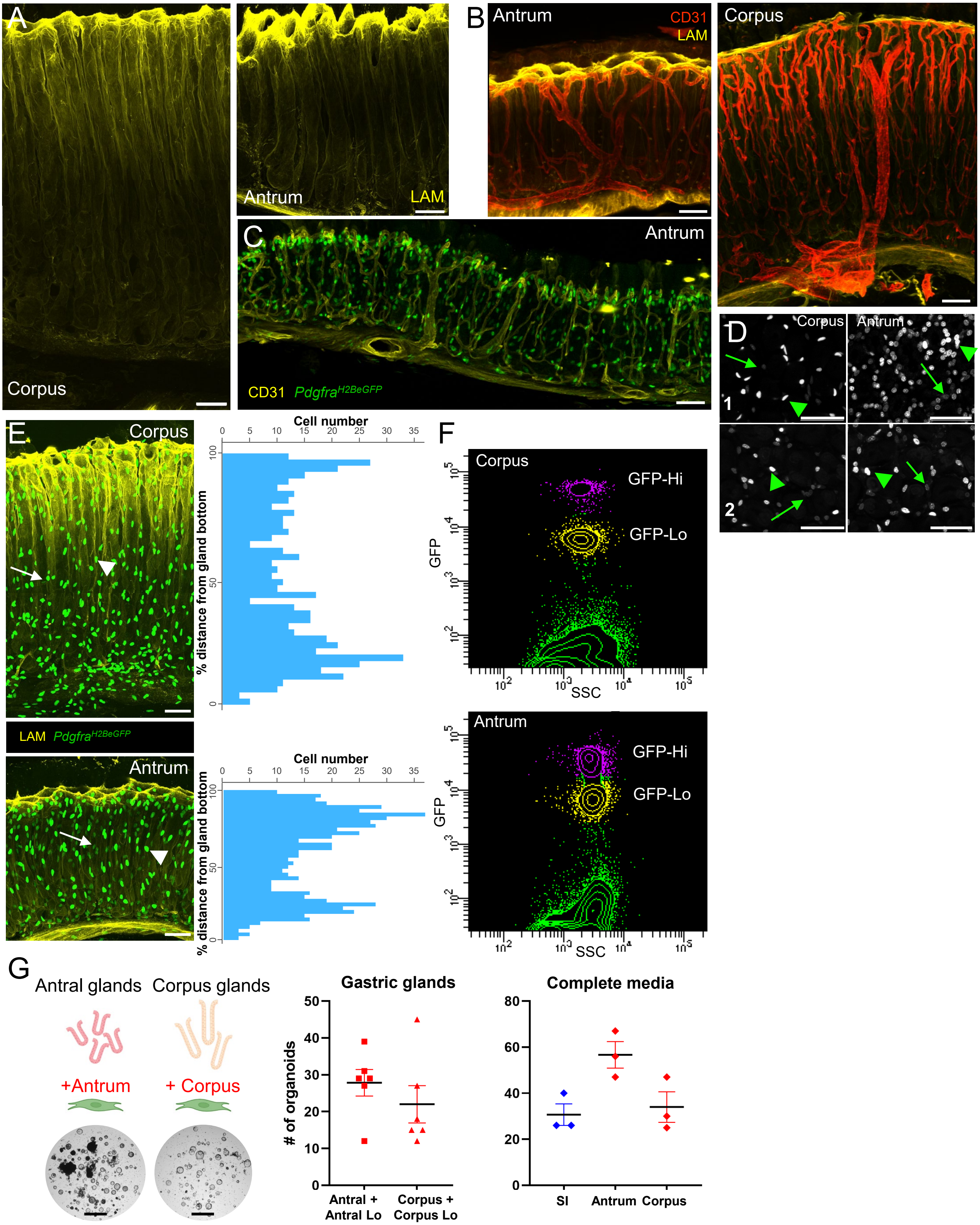

**Figure S1. Structure and organization of gastric corpus and antral mesenchyme.** Related to Figure 1. All scale bars represent 50 µm.

**A)** Whole-mount 3D rendering of Laminin antibody-stained corpus (left) and antral (right) tissue, highlighting the epithelium-mesenchyme interface and gland foveolae (pits).

**B)** Whole-mount 3D rendering of antrum (left) and corpus (right) tissue, showing capillary (CD31, red) dimensions and organization extending from sub-mucosal arterioles. Capillaries branch along the whole length of corpus glands, while antral capillaries branch only near pits. Yellow, Laminin immunostain.

**C)** Whole-mount 3D rendering of $Pdgfra^{H2BeGFP}$ antral glands, showing vascular (CD31, yellow) organization and high density of $\text{PDGFRA}^{\text{Hi}}$ (green) SEMFs across the tissue.

**D)** Grayscale image of GFP signals from the gland cross-sections shown in Figure 1D. These images highlight differential GFP expression in $\text{PDGFRA}^{\text{Hi}}$ (arrowheads) and $\text{PDGFRA}^{\text{Lo}}$ (arrows) cells.

**E)** Whole-mount 3D rendering of $Pdgfra^{H2BeGFP}$ corpus (top) and antral (bottom) glands. Laminin (yellow) marks the basal lamina and $\text{PDGFRA}^{\text{Hi}}$ (arrowheads) and $\text{PDGFRA}^{\text{Lo}}$ (arrows) cells are indicated. Right: GFP signals quantified along gland lengths show highest concentrations at antral pits and the base of corpus glands.

**F)** Separation of $\text{PDGFRA}^{\text{Hi}}$ from $\text{PDGFRA}^{\text{Lo}}$ cells from whole $Pdgfra^{H2BeGFP}$ corpus (top) and antral (bottom) mesenchyme by GFP flow cytometry.

**G)** Left: Matrigel co-culture of glands from different gastric segments with $\text{PDGFRA}^{\text{Lo}}$ cells isolated from the same segment induces organoid growth in the absence of rNOG and RSPO1. Scale bars 400 µm. Right: Quantification of organoid growth and complete media controls (n=3-6). Bars represent mean ±SEM values. Significance of differences determined by one-way ANOVA. Where not indicated, differences were not significant.

Manieri et al, Figure S2

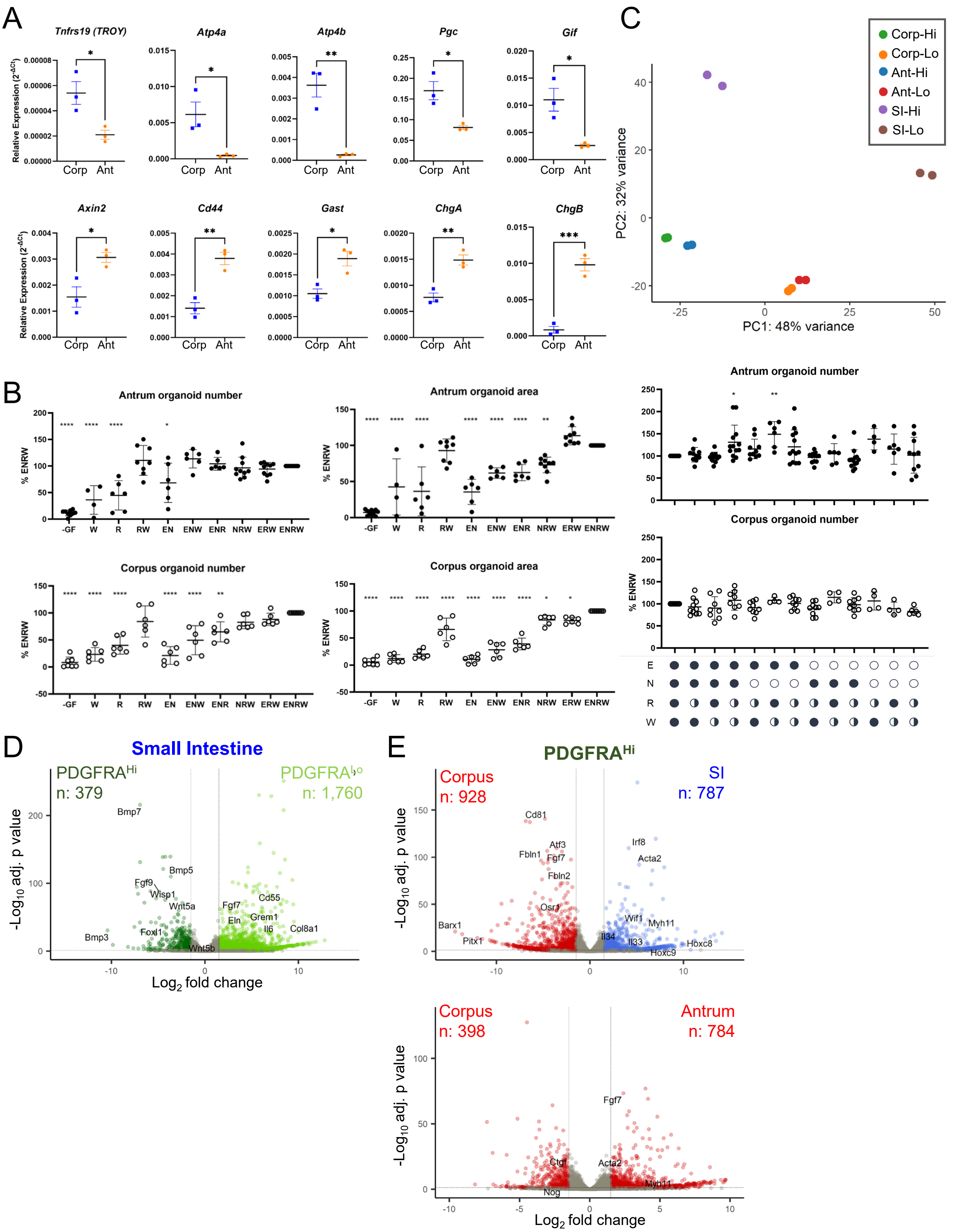

**Figure S2. Properties and dependencies of gastric corpus and antral organoids and differential expression among PDGFRA$^{Hi}$ (SEMFs) and PDGFRA$^{Lo}$ cells.** Related to Figure 2.

**A)** qRT-PCR analysis of corpus and antral epithelial cell markers (relative to *Gapdh*) in organoids cultured for 4 days in complete ENRW medium. Organoids show the expected differences in expression of region-specific marker genes (corpus, blue; antrum, orange).

**B)** Quantitation of organoid growth in different media conditions. -GF, no factors; W, conditioned AFAMIN/WNT3A medium; R, conditioned RSPO1 medium; E, rEGF; N: rNOG. In the right two graphs, circles under the x-axis represent the included factors at full (filled circles) or half concentration or its exclusion (empty circle) from the culture. Bars represent mean ±SEM values (n=4-8 independent experiments). Significance of differences was determined by one-way ANOVA comparing each condition with control organoids grown in ENRW medium at full concentrations. *$p$ <0.05; **$p$< 0.01; ****$p$ <0.0001.

**C)** Principal components analysis (PCA) of duplicate bulk RNA-seq libraries from the indicated cell types, showing high concordance between replicates. SI, small intestine; Corp, corpus; Ant, antrum; Hi: PDGFRA$^{Hi}$ cells (SEMFs); Lo: PDGFRA$^{Lo}$ cells.

**D-E)** Genes differentially expressed ($q$ <0.05; $\log_2$ fold-difference >1.5) between PDGFRA$^{Hi}$ and PDGFRA$^{Lo}$ cells isolated from the SI (**D**), between PDGFRA$^{Hi}$ cells from the gastric corpus and the SI (top, **E**) and between corpus and antrum (bottom, **E**). Selected genes are labeled.

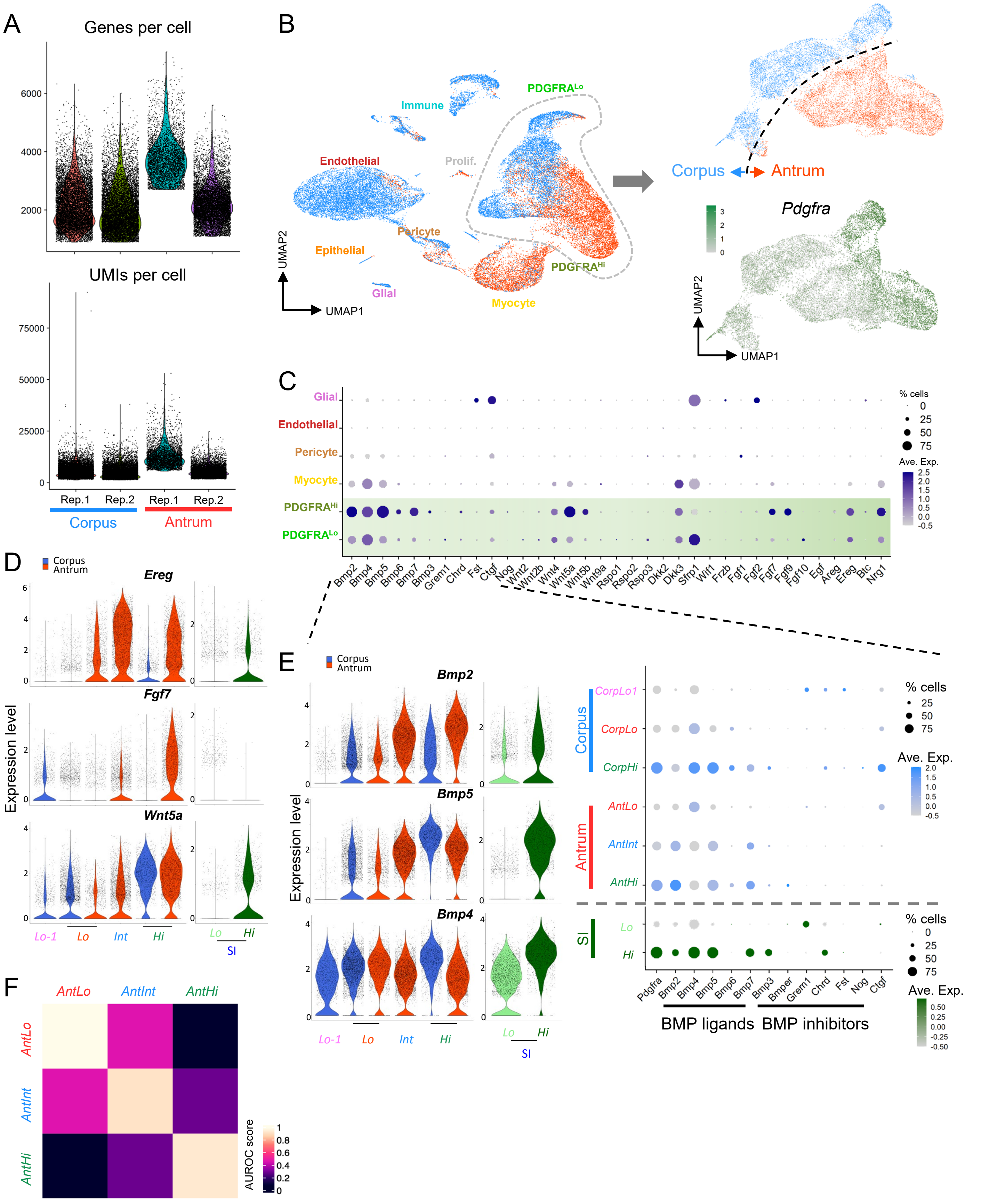
A
Genes per cell
UMIs per cell
Rep.1
Rep.2
Rep.1
Rep.2
Corpus
Antrum
B
Immune
PDGFRA^Lo
Endothelial
Prolif.
Pericyte
Epithelial
PDGFRA^Hi
Glial
Myocyte
UMAP2
UMAP1
Corpus
Antrum
Pdgfra
C
Glial
Endothelial
Pericyte
Myocyte
PDGFRA^Hi
PDGFRA^Lo
% cells
Ave. Exp.
Bmp2 Bmp4 Bmp5 Bmp6 Bmp7 Bmp3 Grem1 Chrd Fst Ctgf Nog Wnt2 Wnt2b Wnt4 Wnt5a Wnt5b Wnt9a Rspo1 Rspo2 Rspo3 Dkk2 Dkk3 Sfrp1 Wif1 Frzb Fgf1 Fgf2 Fgf7 Fgf9 Fgf10 Egf Areg Ereg Btc Nrg1
D
Corpus
Antrum
Ereg
Fgf7
Wnt5a
Expression level
Lo-1
Lo
Int
Hi
Lo
Hi
SI
E
Corpus
Antrum
Bmp2
Bmp5
Bmp4
Expression level
Lo-1
Lo
Int
Hi
Lo
Hi
SI
CorpLo1
CorpLo
CorpHi
AntLo
AntInt
AntHi
Corpus
Antrum
SI
Lo
Hi
% cells
Ave. Exp.
Pdgfra Bmp2 Bmp4 Bmp5 Bmp6 Bmp7 Bmp3 Bmper Grem1 Chrd Fst Nog Ctgf
BMP ligands
BMP inhibitors
F
AntLo
AntInt
AntHi
AUROC score

**Figure S3. Gastric corpus and antral mesenchymal transcripts at single-cell resolution.** Related to Figure 3.

**A)** Parameters of data quality from replicate samples (Rep.) of single mesenchymal cells. UMI: unique molecular identifier.

**B)** Left: Delineation of mesenchymal populations by Uniform manifold approximation and projection (UMAP) of 20,624 corpus (blue dots) and 13,821 antral (orange dots) cells resolved by differential gene expression. mRNA profiles of all cell types, except *Pdgfra*-expressing cells, from the two gastric segments overlap extensively. Right: UMAP plot of these *Pdgfra*$^{+}$ cells extracted from the data on whole mesenchyme separate objectively according to their corpus (6,594 cells) or antral (5,896 cells) origins. Projection of *Pdgfra* transcript density onto the UMAP plot shows *Pdgfra*$^{Hi}$ cells toward the right, *Pdgfra*$^{Lo}$ cells toward the left, and a previously unapparent subset of antral cells with intermediate *Pdgfra* mRNA levels (AntInt, see Figure 3).

**C)** Relative expression of selected BMP, Wnt, FGF, and EGF pathway genes in all mesenchymal populations identified by scRNA-seq. Circle diameters represent the fraction of cells expressing a gene and fill shading, from dark to light purple, represents the normalized average expression in each population. These signaling genes express principally in PDGFRA$^{+}$ cells.

**D)** Relative expression of *Ereg, Fgf7* and *Wnt5a* genes in the identified *Pdgfra*-expressing populations and the corresponding SI cells. Blue, corpus; red, antrum; green, SI.

**E)** Left: Relative expression of *Bmp* ligand genes in the identified *Pdgfra*-expressing populations and the corresponding SI cells. Blue, corpus; red, antrum; green, SI. Right: Relative expression of selected BMP and BMPi genes in gastric and SI PDGFRA$^{+}$ populations. Circle sizes represent the percentage of expressing cells and fill colors represent normalized average expression. Gastric PDGFRA$^{+}$ cells provide a higher BMP tone than in the SI.

**F)** MetaNeighbour AUROC (area under the receiver operating characteristic) analysis shows AntInt population is more similar to AntLo than to AntHi.

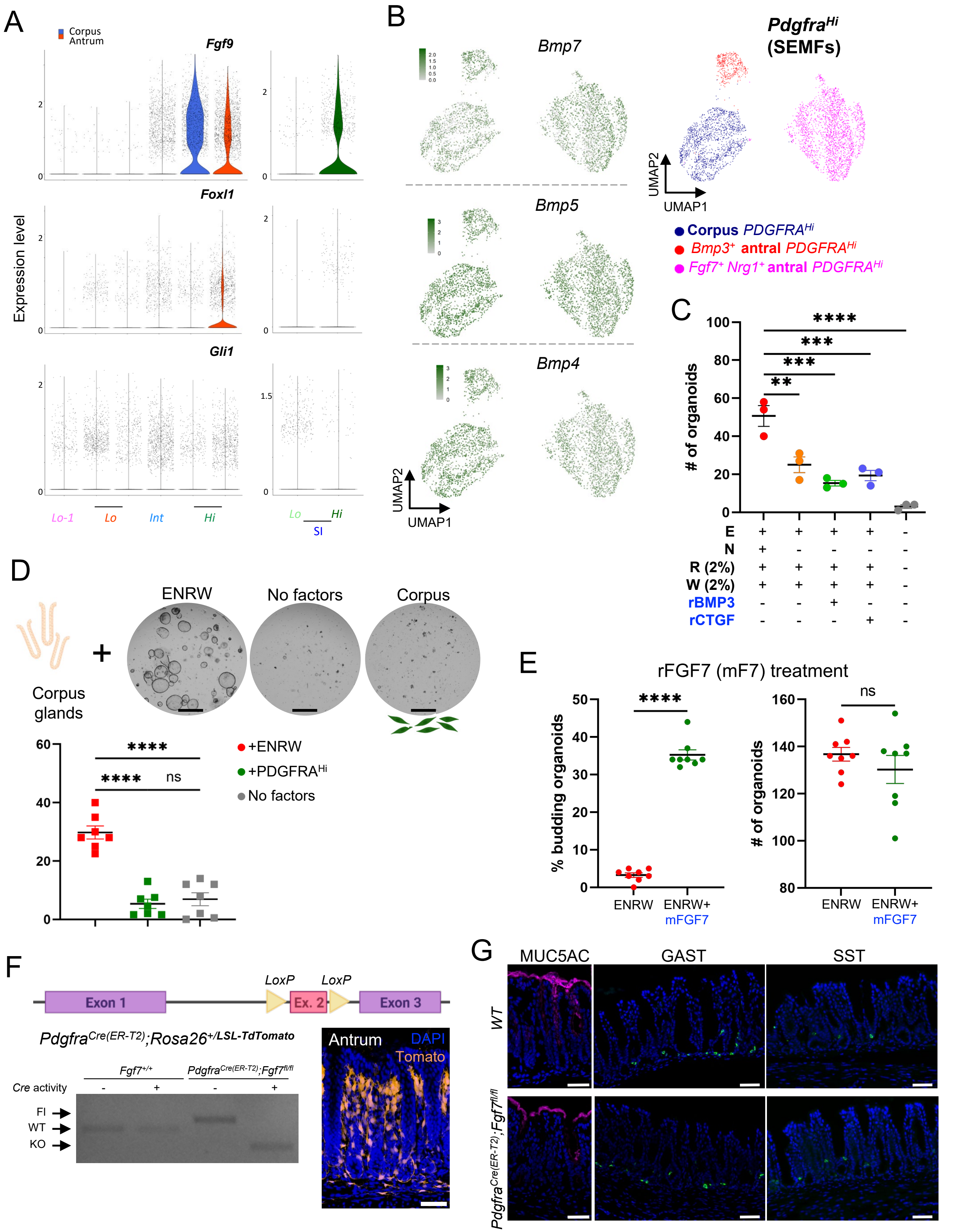
A
Corpus
Antrum
Fgf9
Foxl1
Gli1
Expression level
Lo-1
Lo
Int
Hi
Lo
Hi
SI
B
Bmp7
Bmp5
Bmp4
UMAP2
UMAP1
PdgfraHi
(SEMFs)
Corpus PDGFRAHi
Bmp3+ antral PDGFRAHi
Fgf7+ Nrg1+ antral PDGFRAHi
C
### of organoids
E
N
R (2%)
W (2%)
rBMP3
rCTGF
D
ENRW
No factors
Corpus
Corpus glands
+ENRW
+PDGFRAHi
No factors
ns
E
rFGF7 (mF7) treatment
% budding organoids
### of organoids
ENRW
ENRW+ mFGF7
ns
F
LoxP
LoxP
Exon 1
Ex. 2
Exon 3
PdgfraCre(ER-T2);Rosa26+/LSL-TdTomato
Fgf7+/+
PdgfraCre(ER-T2);Fgf7fl/fl
Cre activity
Fl
WT
KO
Antrum
DAPI
Tomato
G
MUC5AC
GAST
SST
WT
PdgfraCre(ER-T2);Fgf7fl/fl

**Figure S4. Characteristics of regionally distinct gastric SEMF cells.** Related to Figure 4.

**A)** Relative expression of *Fgf9, Foxl1* and *Gli1* genes in the identified *Pdgfra*-expressing populations and the corresponding SI cells. *Fgf9* expression distinguish SEMFs from corpus and SI. Blue, corpus; red, antrum; green, SI.

**B)** UMAP plots of gastric PDGFRA$^{Hi}$ cells extracted from scRNA analysis of whole mesenchyme, resolved into 3 populations: homogeneous corpus cells and two antral cell types, one expressing high *Bmp3* and the other expressing high *Fgf7* and *Nrg1*. Other BMP genes, e.g., *Bmp7, Bmp5* and *Bmp4*, express uniformly among the PDGFRA$^{Hi}$ populations (SEMFs – see also Figure 4B).

**C)** Quantitation of rBMP3 and rCTGF effects on antral organoid formation. Antral glands were treated with these recombinant factors in addition to media with reduced concentrations (2%) of RSPO1 and AFAMIN/Wnt3a and lacking rNOG. Optimized for sensitive detection of BMPi effects, these conditions, failed to elicit such activity from rBMP3 or rCTGF (n= 3 independent experiments). Bars represent mean ±SEM values. Significance of differences was determined by one-way ANOVA. **p <0.01; ***$p$ <0.001; ****$p$ <0.0001.

**D)** Organoid formation from corpus glands co-cultured in Matrigel with PDGFRA$^{Hi}$ cells (SEMFs) isolated from mesenchyme in the same gastric region (n= 7 independent experiments). SEMFs do not induce organoids in the absence of recombinant support factors. Bars represent mean ±SEM values. Significance of differences was determined by one-way ANOVA. ****$p$ <0.0001; ns, not significant. Scale bars 400 µm.

**E)** Quantitation of budding structures and organoid numbers after treatment of antral glands with rFGF7 added to complete ENRW medium. rFGF7 increases budding without affecting organoid numbers (n= 8 independent experiments). Bars represent mean ±SEM values. Significance of differences was determined by t-student statistical analysis. ****$p$ < 0.0001; ns, not significant.

**F)** Schematic representation of an engineered floxed *Fgf7*$^{fl/fl}$ allele. The agarose gel below resolves genotyping PCR products in mesenchymal cells isolated from *Fgf7*$^{+/+}$ (wild-type controls) and *Pdgfra*$^{Cre(ER\text{-}T2)}$*;Fgf7*$^{fl/fl}$ animals with (2 mg tamoxifen injected i.p. on 4 consecutive days) or without activation of Cre recombinase. A representative antral tissue section (n= 3 mice) from a *Pdgfra*$^{Cre(ER\text{-}T2)}$*;Rosa26*$^{LSL\text{-}tdTomato}$ mouse shows abundant Cre activity in PDGFRA$^{Hi}$ cells after tamoxifen exposure (see also Figure 1E and Videos S5 and S6).

**G)** Immunostained antral tissues from wild-type (WT) and *Pdgfra*$^{Cre(ER\text{-}T2)}$*;Fgf7*$^{fl/fl}$ mice reveal no discernible impact of mesenchymal FGF7 deficiency on MUC5AC expression and secretion or on the number and distribution of enteroendocrine GAST- and SST-producing cells. n= 2-3 independent experiments for each marker. Scale bars 50 µm.

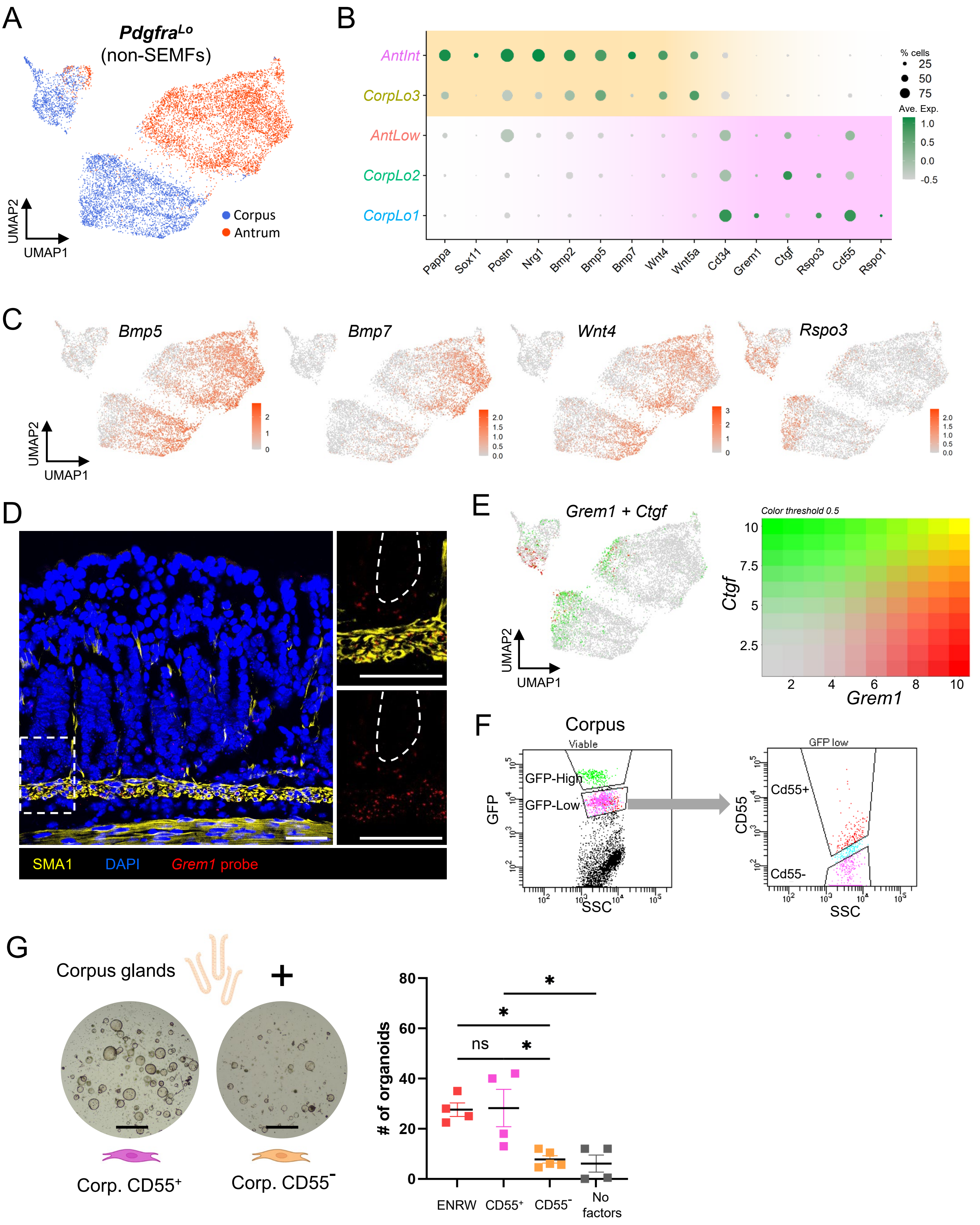
A
PdgfraLo
(non-SEMFs)
UMAP2
UMAP1
Corpus
Antrum
B
AntInt
CorpLo3
AntLow
CorpLo2
CorpLo1
Pappa
Sox11
Postn
Nrg1
Bmp2
Bmp5
Bmp7
Wnt4
Wnt5a
Cd34
Grem1
Ctgf
Rspo3
Cd55
Rspo1
% cells
25
50
75
Ave. Exp.
1.0
0.5
0.0
-0.5
C
Bmp5
Bmp7
Wnt4
Rspo3
UMAP2
UMAP1
D
SMA1
DAPI
Grem1 probe
E
Grem1 + Ctgf
UMAP2
UMAP1
Color threshold 0.5
Ctgf
Grem1
F
Corpus
Viable
GFP-High
GFP-Low
GFP
SSC
GFP low
Cd55+
Cd55-
CD55
SSC
G
Corpus glands
+
Corp. CD55+
Corp. CD55-
### of organoids
ns
*
ENRW
CD55+
CD55-
No factors

**Figure S5. Characteristics and functional activity of regionally distinct gastric PDGFRA$^{Lo}$ and AntInt cells.** Related to Figure 5.

**A)** Resolution of gastric PDGFRA$^{+}$ cells other than SEMFs into populations that segregate largely on the basis of their corpus (blue) or antral (orange-red) origins.

**B)** Relative expression of marker genes in five gastric mesenchymal populations identified by scRNA-seq. Circle diameters represent the fraction of cells expressing a gene and fill colors represent normalized average expression. CorpLo3 shares markers previously identified (see Figure 3F) in AntInt, while other similarities are evident among AntLo, CorpLo1, and CorpLo2.

**C)** *Bmp5, Bmp7, Wnt4*, and *Rspo3* transcript densities projected onto the UMAP plot. CorpLo3 and Ant-Int are the main populations expressing BMP-ligand and non-canonical Wnt genes, while AntLo, Corp1, and Corp2 express *Rspo3*, a potentiator of canonical Wnt signaling.

**D)** In situ hybridization (RNAscope) of antral tissue sections localizes *Grem1* in the muscolaris mucosae and in the area corresponding to PDGFRA$^{Lo}$ cells near the epithelial stem cell zone. Images represent fields examined in 1 experiments. Scale bars 50 μm.

**E)** *Grem1*, *Ctgf*, and their co-expression densities projected onto the UMAP plot. Both genes are expressed, but rarely together, in the AntLo, Corp1, and Corp2 clusters.

**F)** Flow cytometry for GFP separates whole corpus mesenchyme into PDGFRA$^{Hi}$, PDGFRA$^{Lo}$, and PDGFRA$^{-}$ cells (top) and additional staining with CD55 antibody separates PDGFRA$^{Lo}$ cells into CD55$^{+}$ and CD55$^{-}$ fractions (bottom). Isolation of the corresponding antral population is shown in Figure 5F.

**G)** Co-culture of corpus glands with CD55$^{+}$ PDGFRA$^{Lo}$ or CD55$^{-}$ PDGFRA$^{Lo}$ cells. As observed for the antrum, the CD55$^{+}$ but not the CD55$^{-}$ fraction induced organoids in the absence of any recombinant factors (n= 4 independent experiments). **p* <0.05; ns, not significant. Where not indicated, differences were not significant. Scale bars 400 μm.
